# Structural Insights into Kainate Receptor Desensitization

**DOI:** 10.1101/2025.03.27.645769

**Authors:** Changping Zhou, Guadalupe Segura-Covarrubias, Nami Tajima

**Affiliations:** Department of Physiology and Biophysics, Case Western Reserve University School of Medicine, Ohio, 44106, USA

**Keywords:** Kainate-type ionotropic glutamate receptors, structure, desensitization mechanism

## Abstract

Kainate receptors (KARs), along with AMPA and NMDA receptors, belong to the ionotropic glutamate receptor (iGluR) family and play critical roles in mediating excitatory neurotransmission throughout the central nervous system. KARs also regulate neurotransmitter release and modulate neuronal excitability and plasticity. Receptor desensitization plays a critical role in modulating the strength of synaptic transmission and synaptic plasticity. While KARs share overall structural similarity with AMPA receptors, the desensitized state of KARs differs strikingly from that of other iGluRs. Despite extensive studies on KARs, a fundamental question remains unsolved: why do KARs require large conformational changes upon desensitization, unlike other iGluRs? To address this, we present cryo-electron microscopy structures of GluK2 with double cysteine mutations in non-desensitized, shallow-desensitized and deep-desensitized conformations. In the shallow-desensitized conformation, two cysteine crosslinks stabilize the receptors in a conformation that resembles the desensitized state of AMPA receptors. However, unlike the tightly closed pore observed in the deep-desensitized KAR and desensitized AMPAR conformations, the channel pore in the shallow-desensitized state remains incompletely closed. Patch-clamp recordings and fluctuation analysis suggest that this state remains ion-permeable, indicating that the lateral rotational movement of KAR ligand-binding domains (LBDs) is critical for complete channel closure and stabilization of the receptor in desensitization states. Together with the multiple conformations representing different degree of desensitization, our results define the unique mechanism and conformational dynamics of KAR desensitization.

**Highlights:**

1. We present cryo-EM structures of GluK2 kainate receptors with engineered cysteine crosslinks at the inter-dimer interface, which restrict subunit lateral rotation and attenuate receptor desensitization.
2. The structure of GluK2 double cysteine mutant in complex with the allosteric potentiator BPAM344 and glutamate represents a non-desensitized state, highlighting the critical conformational changes required for ion channel gating.
3. The glutamate-bound GluK2 mutant adopts multiple conformations, representing both shallow- and deep-desensitized states. Electrophysiological recordings indicate that the GluK2 kainate receptor mutant recovers from desensitization more rapidly, resembling AMPA receptors. Our structural and functional data suggest that shallow-desensitized KARs remain conductive, implying that the large lateral LBD rotation during KAR desensitization is essential for complete channel closure, distinguishing KARs from other iGluRs.

---

Kainate receptors (KARs) belongs to the ionotropic glutamate receptor (iGluR) superfamily together with α-amino-3-hydroxy-5-methyl-4-isoxazolepropionic acid (AMPA), *N*-methyl-d-aspartate (NMDA) and delta receptors. They are cation selective ligand-gated ion channels activated by an excitatory neurotransmitter glutamate and mediate fast excitatory synaptic transmission in mammalian brains^1,2^. In addition to their role in mediating the synaptic transmission at post-synapses, KARs also function pre- and extra-synaptically, and regulate excitatory and inhibitory neurotransmitter release and modulate synaptic excitability^3–7^. Abnormal expression and dysfunction of KARs are associated with various neurological diseases and disorders, including schizophrenia, mood disorders, epilepsy, and Huntington’s disease^8–17^. Therefore, KARs are considered potential therapeutic drug targets due to their physiological and pathophysiological roles^18–21^.

Kainate receptors (KARs) assemble as homo- or heterotetramers composed of five subunits: GluK1–5^22–25^. Extensive studies revealed that KARs adopt a Y-shaped architecture with four distinct layers: the amino-terminal domain (ATD), which is critical for receptor assembly; the ligand-binding domain (LBD), which contains ligand-binding sites and regulates channel gating; the transmembrane domain (TMD), which forms the ion channel within the plasma membrane; and a short disordered C-terminal domain (CTD), which interacts with intracellular proteins^1,26^. Recently determined structures of intact KARs in various liganded states have elucidated the structure-function correlations underlying the mechanisms of activation/desensitization^25,27–31^, allosteric modulation^32^, inhibition^33^, and channel block^34^. A characteristic structural feature of KARs lies in their unique conformation during desensitization. Structures of intact GluK2, first reported in 2014 by the Mayer/Subramaniam lab, have indicated that KAR LBD dimer assemblies are completely dissociated upon desensitization, with a large horizontal-plane rigid-body rotation of the LBDs in the BD subunits, resulting in an approximately four-fold symmetric structures, while other iGluRs typically retain two-fold symmetric conformations viewed from the top when they are desensitized^26,28^. In addition, a key functional characteristic of KARs is their significantly slower recovery from desensitization — over 50 fold slower compared to AMPARs^35,36^— which limits KAR mediated rapid and high-frequency transmission in neurons^37^. Due to the rapid onset of desensitization induced by the neurotransmitter glutamate, which closes the ion channel within a few milliseconds, previous KAR structures in complex with glutamate have been stabilized in fully or deep-desensitized states, with the ion channel fully closed^25,27–31,33,38–40^. Despite their unique structural and functional features, why only KARs exhibit such a strikingly different conformation, which is not conserved in other iGluRs, and how these drastic conformational changes control receptor gating remains elusive.

To address this outstanding question, we present the cryo-electron microscopy (cryo-EM) structure of the GluK2 KAR in multiple functional states, including non-desensitized, shallow-desensitized, intermediate (between shallow and deep desensitized), and deep-desensitized conformations, using cysteine cross-linking in the presence of glutamate and the positive allosteric modulator BPAM344. The structural comparison revealed that the large in-plane rotation of LBDs in both BD subunits and the resulting twisted motion of the LBD-TM3 linkers are essential for tight ion channel closure and stabilization of receptors in the deep-desensitized state—a feature not observed in AMPARs and NMDARs. Functionally, the crosslinked GluK2 mutant exhibits large fractional steady-state currents, as revealed by patch-clamp recording. The stationary noise analysis results indicate that the GluK2 mutant remains conductive when desensitized, resembling the behavior of the conductive GluA2(R) AMPAR under similar conditions^41^. These structural and functional results show that robust in-plane rotations of the LBDs are essential for proper ion channel closure. Overall, our study provides new insights into the unique desensitization mechanism of the KAR subfamily.

## Results

### Trapping GluK2 KARs in a non-desensitizing state

Previous studies have shown that LBD dimer reorientations switch the gating of KARs^27–29^. To gain insight into the detailed conformational transitions from active to desensitized states, we first generated a homology model of the intact open GluK2 receptor, comprising the ATD, LBD, and TMD layers based on the structure of the activated GluA2 AMPA receptor in the presence of glutamate, potentiator, and auxiliary subunit in its higher conductive open state (Protein Data Bank (PDB) code 5WEO)^42–44^ using SWISS-MODEL^45–49^ (Fig. S1A, B). In the model, the G/E helices of subunit A and the K helix of subunit B come into close proximity (Fig. S1C, D), resembling the recently determined structure of GluK2 bound to glutamate, an allosteric positive modulator BPAM344, and Concanavalin A (ConA) in an open state, using time-resolved cryo-EM (PDB code 9B36)^27^. Specifically, the distances between the Cα atoms of K676 in subunit A and N802 in subunit B are 6.2 Å and 6.7 Å in the homology model and the active/open GluK2 structure, respectively. By contrast, these distances are much greater in the BAPM344-bound (no agonist supplemented) inactive state (PDB code 8FWS)^32^, as well as in the antagonized (PDB code 5KUH)^28^, and desensitized (PDB code 9B38, 5KUF)^27,28^ states (Fig. S1D). Cysteine crosslinking approaches have been used to analyze the dynamics and the channel function of iGluRs^50–54^. Additionally, a recent report showed that the conformational changes of KARs upon activation resembles those of AMPARs^27^. Based on these reports, we hypothesized that an engineered disulfide bond between residues K676 and N802 would restrict the mobility of the LBD and trap a dimer-of-dimers LBD conformation, potentially altering desensitization kinetics. To test this hypothesis, we introduced a pair of cysteine residues designed to form disulfide bonds between the LBD dimer pairs (Fig. S1 E, F) and assessed the channel activity of the full-length rat GluK2 K676C/N802C mutant using whole-cell patch-clamp recordings. Note that the distance between the Cα atoms of two cysteines required to form a disulfide bond is reported to be less than 7 Å^55^.

GluK2 wild type (WT) demonstrated the rapid desensitization, as observed previously^35^. By contrast, the GluK2 K676C/N802C mutant, activated by 1 mM glutamate, showed earlier rapid desensitization and a steady-state current (Fig. 1A). When we assessed the GluK2 K676C and GluK2 N802C single cysteine mutants, both exhibited nearly complete desensitization, similar to GluK2 WT, without displaying the steady-state current observed when the GluK2 K676C/N802C double cysteine mutant was activated by glutamate (Fig. S2A). These results confirm that disulfide bond formation between K676C and N802C are responsible for generating the steady-state current.

**Figure 1.**
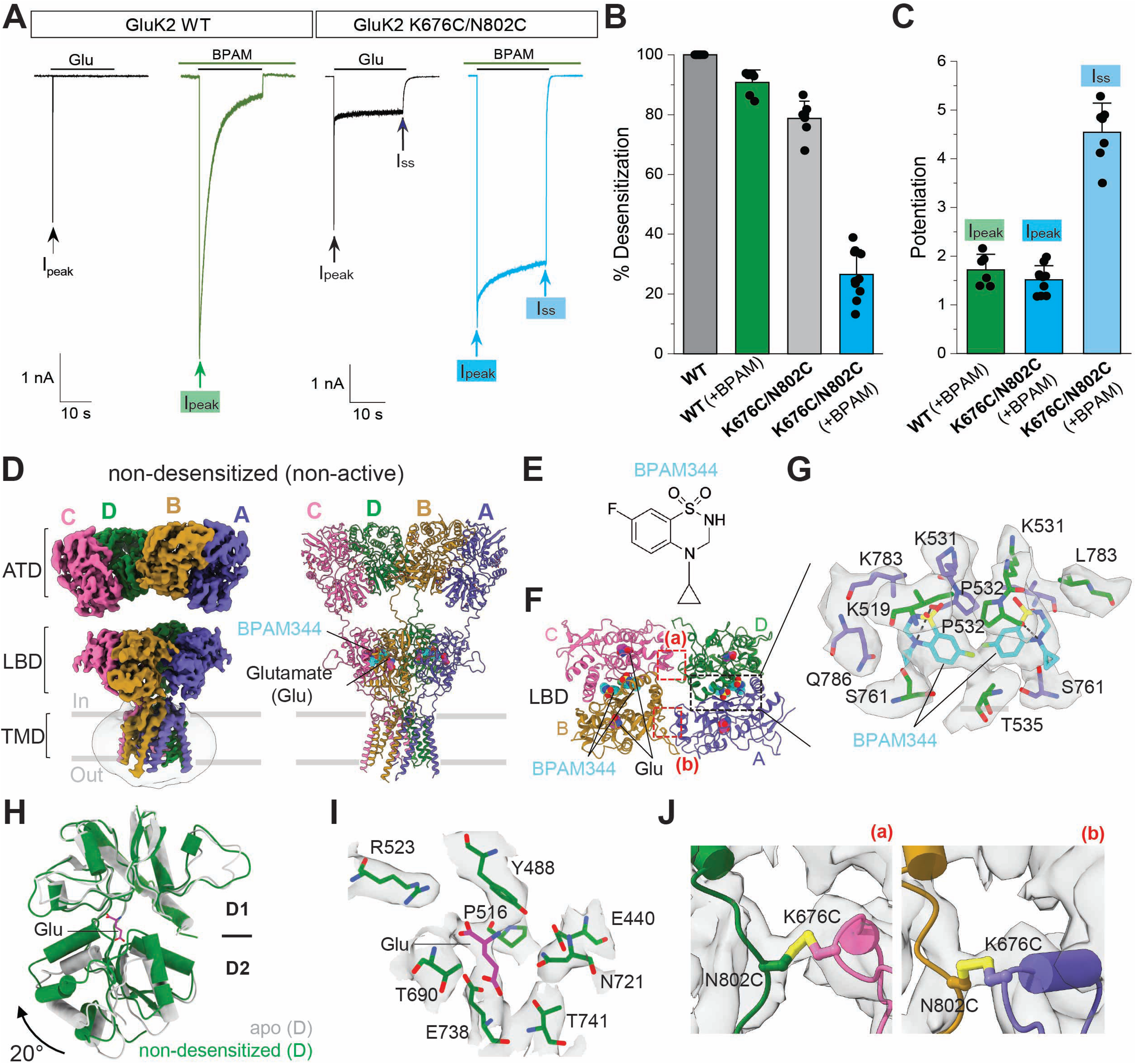
Functional and structural characterization of GluK2 K676C/N802C. **(A)** Representative whole-cell patch-clamp trace showing response of GluK2 WT or GluK2 K676C/N802C to a 20-second application of 1 mM glutamate in the absence and presence of BPAM344 (BPAM). Arrows indicate the maximum peak current (I_peak_) and the steady-state current (I_ss_). **(B)** Quantification of percent desensitization for currents evoked by 1 mM glutamate. **(C)** Potentiation of the peak current amplitudes (I_peak_) of GluK2 WT (left), the peak current amplitudes (I_peak_) of K676C/N802C (middle), and the steady-state current amplitude (I_ss_) of GluK2 K676C/N802C measured in the presence of BPAM344 compared to currents in the absence of BPAM344. **(D)** Cryo-EM map and model of GluK2 K676C/N802C in complex with glutamate and BPAM344 in a non-desensitized/non-active state. **(E)** Chemical structure of BPAM344. **(F)** Structure of the LBD layer viewed parallel to the membrane, with BPAM344 and glutamate shown as space-filling models, colored pink and cyan, respectively. **(G)** Close-up view of the BPAM344 binding site at the AD subunit interface, indicating by a black dotted box in panel F. **(H)** Comparison of the apo GluK2 LBD (PDB code 9CAZ) and the glutamate-bound GluK2 LBD in the non-desensitized LBD conformation. The LBD is a bi-lobed structure, consisting of D1 and D2 lobes, with glutamate binding in the cleft between them. The D1 lobe is superimposed, and the rotation angles to align the D2 residues were calculated. The glutamate-bound bi-lobe is closed 20° than the apo state LBD conformation. **(I)** Cryo-EM density for bound glutamate and the surrounding residues in the agonist binding pocket. **(J)** The cryo-EM densities for disulfide bonds between K676C and N802C in AB and CD subunits. The crosslinks in the LBD layers are indicated by the red dotted box in the panel F.

An earlier study showed that BPAM344 significantly slows the desensitization kinetics of GluK2 WT and enhances the glutamate-evoked current by 140-fold and 21-fold, respectively^56^. To assess how BPAM344 modulates the activation and desensitization of the GluK2 K676C/N802C mutant, we next analyzed its effect. Indeed, the percent desensitization of GluK2 K676C/N802C elicited by 1 mM glutamate is dramatically decreased in the presence of 0.5 mM BPAM344, with only 25% desensitization observed after 20 seconds (Fig. 1 A, B). While the peak current amplitude (I_peak_) was increased 1.5-fold (1.5 ± 0.3, n=10), similar to the WT in the presence of BPAM344, the steady-state current (I_ss_) amplitudes were also significantly increased, by approximately 4.7-fold (4.7 ± 0.6, n=10) (Fig. 1C). Thus, we sought to capture the conformation of receptors in this non-desensitizing state using cryo-EM.

### GluK2 K676C/N802C adopts multiple conformations

We expressed the intact homotetrameric GluK2 KAR incorporating K676C/N802C double cysteine mutations in HEK293S GnTl^-^ cells and purified the proteins in the detergent, n-Dodecyl-B-D-Maltoside (DDM). The final protein sample (Fig. S2B, C) was pre-incubated with 1 mM glutamate, with or without 0.5 mM BPAM344, prior to grid freezing. We initially performed 3-dimensional (3D) refinement without imposing symmetry (Table S1). Our initial global reconstruction of the glutamate/BPAM344-supplemented dataset generated six main classes (Fig. S3a). One of the classes has a two-fold rotational symmetrical conformation (Fig. 1D), with the extracellular domains resembling the open-state conformation (PDB code 9B36)^27^. To improve the density, we further performed signal subtraction on the ATD and LBD-TMD layers separately. The local refinements with focused masks yielded 3.7 Å and 3.9 Å reconstruction of the ATD and LBD-TMD, respectively, enabling reliable model building (Fig. S3a, S4 A–C). The reconstruction of the LBD-TMD showed clear density for BPAM344 at the LBD dimer interface consistent with previous reports^27,32,57^ (Fig. 1E–G). All four LBD bi-lobes are closed in the presence of glutamate, compared to its open conformation in an apo GluK2 (PBD code 9CAZ) structure (Fig. 1H, I). While the extracellular domain adopts an active-like conformation, the ion channel pore remains closed. Therefore, we refer to this class as a non-desensitized conformation.

In contrast, the remaining five classes showed no BPAM344 molecules at the LBD D1-D1 interface, leading to D1-D1 rupture. In all of these five classes, the LBD bi-lobes of all four subunits are tightly closed in the presence of glutamate, with closure degrees ranging from 23° to 27°, indicating glutamate binding to the receptors. The overall structures of these five conformations are stabilized in two- or four-fold symmetrical, or asymmetrical arrangements. In order to analyze conformations in the absence of allosteric potentiators, we also collected cryo-EM data of GluK2 K676C/N802C supplemented with glutamate. These data revealed four main three-dimensional (3D) classes (Fig. S3b), each are superimposable with the glutamate-bound, BPAM344-unbound classes from the BPAM344-supplemented sample (Fig. S3a). Thus, we combined these two data sets and further performed processing and refinement to improve the density quality. We defined these classes as shallow-desensitized, intermediate, and deep-desensitized states, based on the observations described below. Ultimately, the final resolutions of desensitized classes range from 3.7 Å to 4.0 Å (Fig. S4).

### GluK2 K676C/N802C mutant structure bound to glutamate and BPAM344 represents a non-desensitized conformation

While all six 3D reconstructions showed three domains—ATD, LBD, and TMD—as previously reported^25,28,30,31,33^, each class showed distinct conformations (Fig. S3). In the non-desensitized structure, two molecules of BPAM344 per LBD dimer bind and stabilize the LBD D1-D1 interfaces, with each BPAM344 forming hydrogen bonds with the side chain of P532, as recently reported^27,32,57^ (Fig. 1G). In addition to BPAM344, clear densities corresponding to disulfide bonds between K676C and N802C were observed in this conformation (Fig. 1J). These two disulfide bonds trapped the LBDs in a dimer-of-dimers conformation in the presence of the agonist, as designed (Fig. 1D, F). Previously, the structure of open GluK2 was determined in the presence of two positive allosteric modulators, concanavalin A (ConA), BPAM344, as well as the agonist glutamate (PDB code 9B36)^27^. In the structure, the ∼50 kDa ConA dimer, along with BPAM344 and glutamate, binds at the ATD-LBD interface, elevating the ATD by 16 Å and rotating the LBD relative to the ATD layer by 28° (Fig. S5). The same tendency was observed in the presence of NETO auxiliary proteins, which also bind to the ATD and LBD layers and modulate KAR kinetics^30^. Interestingly, a comparison between structures without an orthosteric ligand (PDB codes 8FWQ, 9CAZ)^32^ and our non-desensitized structure revealed that the distance between the center of mass (COM) of the ATD and LBD was also increased by 7.5 Å in the non-desensitized conformation, even in the absence of ConA and NETO auxiliary proteins. By contrast, in the desensitized state, this distance between the ATD and LBD decreased by 2–4 Å compared to the apo state. While, the ATDs in all structures are superimposable and do not undergo any conformational changes (Fig. S5B), the ATD in the non-desensitized class also displayed a rigid-body rotation of 45° counterclockwise (Fig. S5C), following the same movement tendency observed in the open state of KARs^27^ and AMPARs^58^.

Focusing on the LBD layer, the LBDs are stabilized in a dimer-of-dimers arrangement that resembles the open conformation (Fig. 2A), with the D1–D1 association stabilized by BPAM344 and the D2–D2 separation maintained within each LBD dimer in the presence of glutamate (Fig. 2B). Analysis of the center-of-mass (COM) distances between D1-D1 and D2-D2 in the two structures further indicates their similar conformational arrangement. Specifically, the distance between the Cα atoms of S670, located at the bottom of the D2 lobes in the AD and BC subunit pairs, is equally separated as seen in the open-state conformation. Key features of the open-state conformation of iGluRs include increased tension in LBD-TM3 linkers and the expansion of the ion channel pore held by the linkers, particularly at the gate region^1,59^. Thus, we next compared the LBD-TM3 linker arrangement, TM3 helix orientation, and channel openings in the BPAM344-bound (without an orthosteric ligand), non-desensitized, and open GluK2 conformations. The densities of all TM3 helices are well resolved (Fig. S6). In the non-desensitized conformation, the TM3 helices are unwound at A656 in the BD subunits and at E662 in the AC subunits (position (3) and (2’), respectively), and therefore, T660 (BD subunits) and M664 (AC subunits) are pulled away from the ion channel central axis, while BPAM344-bound GluK2 (PDB code 8FWS)^32^ displays an additional half-turn of the helix at the pore entrance, and the TM3 helices unwind at T660 in the AC subunits and at M664 in the BD subunits (position (2) and (1), respectively) (Fig. 2C). Consequently, the top of the gating region, which contains the conserved SYTANLAAF motif^60,61^, is more dilated in the non-desensitized conformation compared to the more tightly closed BPAM344-bound GluK2 conformation. By contrast, the TM3 helices in the open GluK2 conformation (PDB code 9B36)^27^ display kinking at a deeper position, specifically at L655 (position (4)) in all four subunits (Fig. 2D). Additionally, the open GluK2 conformation exhibits the rotational movements of TM3, which put away T652 from the central axis of the pore, thus opening the channel. However, these rotational movements are not observed in the non-desensitized conformation. Overall, the ion channel in the non-desensitized conformation remains closed (Fig. 2E, F).

**Figure 2.**
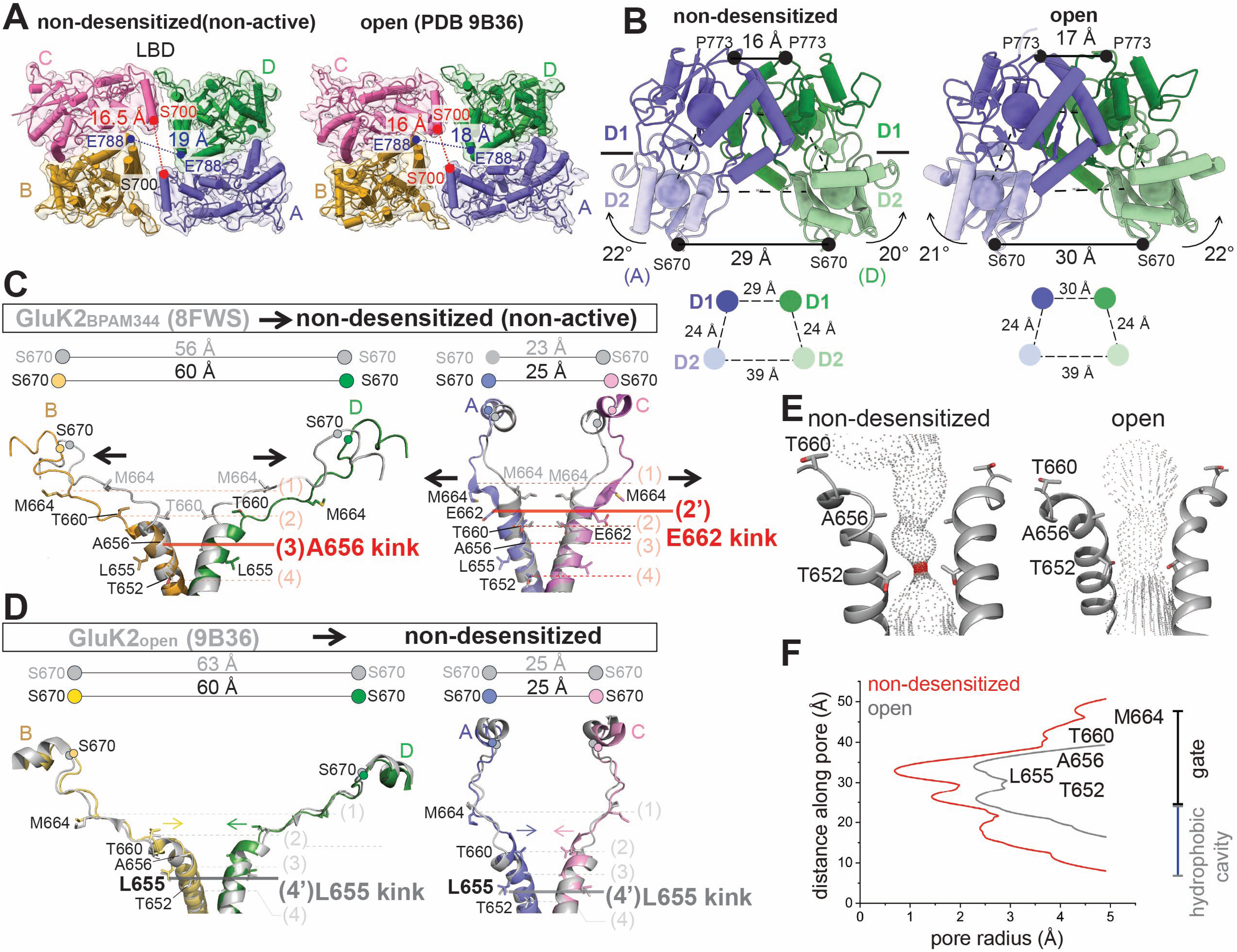
Structural mechanism of GluK2 upon activation and desensitization. **(A)** Comparison of Cryo-EM reconstructions and models of BPAM344- and glutamate-bound GluK2 K676C/N802C in a non-desensitized conformation, and BPAM344-, glutamate-, and Concanavalin A-bound GluK2 in an open (active) state (PDB code 9B36). The distances between S700 in the AC subunits and E788 in the BD subunits at the LBD dimer-dimer interface are colored red and blue, respectively. **(B)** Structural comparison of LBD dimers in the non-desensitized and open GluK2, viewed perpendicular to the membrane. The locations of the centers of mass (COM) of D1 and D2 are indicated by blue and green spheres, respectively, with the COM distances between D1 and D2 of subunit AD LBDs (shown as circles) displayed below. Arrows indicate LBD bi-lobe closure, with the D2 lobe moving closer to D1. Distances between the Cα atoms of P773 (D1-D1 distance) and S670 (D2-D2 distance) in the AD subunits are also indicated. **(C)** Comparison of LBD-TM3 linkers and TM3 formation in BPAM344-bound, agonist-unbound GluK2 (PDB code 8FWS)^32^ and non-desensitized GluK2, showing the subunit BD pair (left) and the subunit AC pair (right). S670 is shown as spheres in corresponding colors for the non-desensitized state and in gray for BPAM344-bound GluK2. The cross-dimer distances between the Cα atoms of S670 (in the BD and AC subunit pairs) are indicated above. Residues forming the ion channel pore are displayed as sticks. The locations of the TM3 gating hinge at A656 (subunit BD) and E662 (subunit AC) in the non-desensitized conformation are highlighted in red. The dotted line indicates the positions of pore-lining residues: (1) M664, (2) T660, (3) A656, and (4) T652. (2’) E662, which serves as the gating hinge, is also highlighted in this panel. Helices surrounding T664 and M664 in subunit BD, as well as helices around M664 in subunit AC, are unwound in the non-desensitized conformation. As a result, the ion channel pore above the gating region in the non-desensitized conformation is wider compared to that in the BPAM344-bound GluK2 structure, with black allows indicating the conformational changes. **(D)** Comparison of LBD-TM3 linkers and TM3 formation in non-desensitized GluK2 and BPAM344-, ConA- and glutamate bound open GluK2 (PDB code 9B36)^27^, showing the subunit BD pair (left) and subunit AC pair (right). The cross-dimer distances between the Cα atoms of S670 are indicated above. Residues forming the ion channel pore are displayed as sticks. The locations of the gating hinge at L655 in A-D subunits of the open GluK2 (PDB 9B36) structure are highlighted in gray, for comparison with the gating hinge in the non-desensitized conformation shown in panel C. The dotted line indicates the positions of pore-lining residues: (1) M664, (2) T660, (3) A656, and (4) T652, as well as (4’) L655, which forms the gating hinge. Yellow, green, blue, and pink arrows indicate the rearrangement of TM3 in each subunit during the transition from the non-desensitized to the open conformation. **(E)** Pore profile in the non-desensitized and open (PBD 9B36) structures. Pore-delineating dots are colored according to pore radius: red for regions with a radius less than 1.1 Å and gray for regions with a radius greater than 1.1 Å, which would allow passage of a dehydrated ion (1.1 Å for calcium). **(F)** The pore radius for the non-desensitized (red) and open (PDB 9B36, gray) conformations, calculated using HOLE.

### Disulfide bonds between K676C-N802C stabilized GluK2 KARs in a desensitized AMPA receptor like conformation

The glutamate-bound, potentiator-unbound GluK2 K676C/N802C generated five classes (Fig. 3A). The LBD layer conformations in these classes appear desensitized; however, they represent different levels of desensitization (Fig. 3B). In one of these desensitized classes, the absence of BPAM344 results in D1–D1 dissociation, a hallmark structural feature of iGluR desensitization, while two disulfide bonds between K676C and N802C (Fig. 3C) stabilize LBD dimers forming a dimer-of-dimers conformation. We refer to this as the ‘shallow-desensitized’ conformation to distinguish it from the typical deep-desensitized conformations previously observed^28^. Another class, termed the intermediate state, contains an LBD dimer on one side and a disrupted dimer on the other. In this class, one LBD dimer pair maintains its dimer formation with D1–D1 dissociation, resembling the shallow-desensitized state, while the disrupted dimer adopts a conformation characteristic of deep-desensitized classes. Therefore, we propose that this structure represents an intermediate state between the shallow- and deep-desensitized conformations in the presence of glutamate. In contrast, the other three structures lacked inter-subunit disulfide bonds between K676C and N802C, resulting in the LBD layers stabilizing an approximately four-fold symmetric conformation, though with varying in-plane rotation angles of the LBDs (Fig. 3B). We observed the formation of a ‘desensitization ring’ (Fig. 3D, E) in all three deep-desensitized conformations, consistent with previous reports^28,29,62^. This ring formation plays a key role in regulating desensitization and recovery from desensitization^28,29^. Accordingly, we classified these classes as ‘deep-desensitized’ states.”

**Figure 3.**
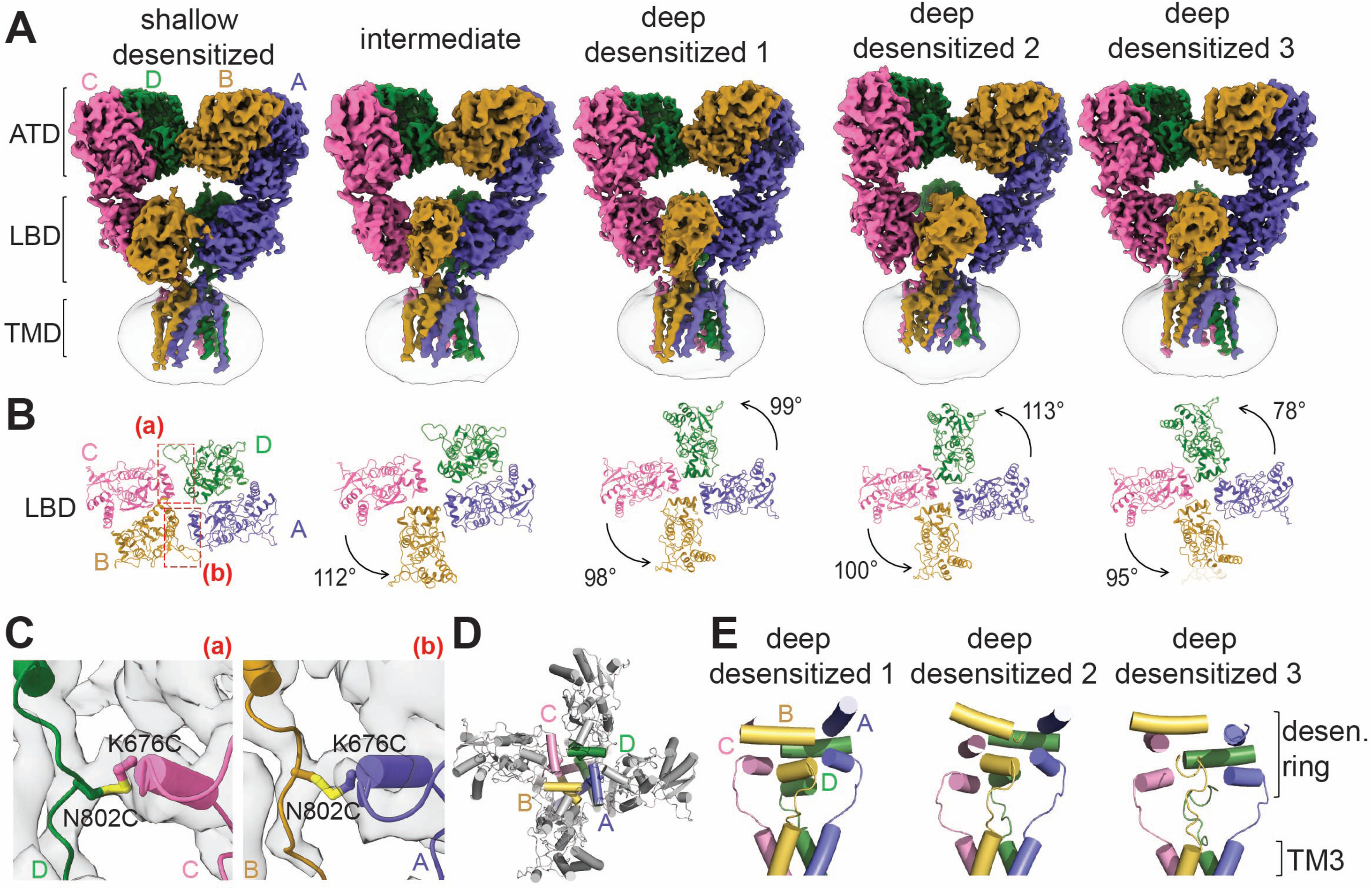
Structures of GluK2 in desensitized states. **(A)** Three-dimensional (3D) cryo-EM reconstructions of glutamate-bound GluK2 K676C/N802C in desensitized conformations. **(B)** Top views of glutamate-bound GluK2 K676C/N802C, focusing on the LBD layers. The in-plane rotation angles of the LBDs are indicated. Disulfide bonds formed between the two LBD dimers are highlighted in red dotted boxes. **(C)** Close-up view of the disulfide bonds between K676C and N802C at the CD subunit interface (left) and the AC subunit interface (right) in the shallow-desensitized conformation. **(D)** Top view of the LBD layer in the deep-desensitized conformation, highlighting helices E and G, which form the desensitization ring. The view is parallel to the membrane, with coloring consistent with panel A. **(E)** The desensitization ring formed in three deep-desensitized conformations, viewed perpendicular to the membrane. The in-plane rotation of the LBD shown in panel B shifts the LBD-TM3 linker, positioning it above the ion channel pore formed by the TM3 helices.

Comparison of the non-desensitized and shallow-desensitized conformations reveals multiple structural rearrangements in the context of the full-length receptor structure. The distance between the ATD and LBD decreases in the shallow-desensitized state, measuring 2–3 Å shorter than in the apo and BPAM-bound GluK2 structures, making it more similar to the deep-desensitized state than to the open-state conformation (Fig. S5A). Focusing on the LBD layer, the LBDs in the AC subunits are rotated by 5° parallel to the membrane when viewed from the top, compared to the non-desensitized conformation (Fig. 4A). Viewed from the side, the entire LBDs appear rolled down (Fig. 4B), which is associated with the rearrangement of the dimer-of-dimers formation and the rupture of the D1-D1 interface. These conformational changes are also observed during AMPAR desensitization^44,63–66^. Therefore, we next compared the shallow-desensitized conformation of GluK2 K676C/N802C with the desensitized conformation of GluA2 AMPAR (PDB code 7RYZ)^64^. Viewed from the top, the distance between the Cα atoms of S700 (S662 in GluA2 AMPAR) in both structures are reduced compared to their respective open states, although the arrangements in these two structures differ due to the restricted mobility of GluK2 caused by cysteine crosslinks (Fig. 4C). The D1 lobes of GluK2 roll down approximately 25°, in contrast to the 33° rolling-down motion observed in the D1 lobes of GluA2 (Fig. 4D). The D2 distances in the shallow-desensitized GluK2 and desensitized GluA2 are comparable, measuring 20 Å and 24 Å, respectively. Overall, the shallow-desensitized conformation of the GluK2 mutant appears to resemble the desensitized conformation of AMPA receptors. However, this conformation in KAR is energetically unstable and has not been observed in the absence of crosslinks. Comparison of the TM3 in the non-desensitized and shallow-desensitized conformations revealed that the pore in the shallow-desensitized state is occluded at T660, and M664 in the BD and AC subunits, respectively (position (2) and (1)) (Fig. 4E), while the ion channel pore in the deep-desensitized conformations is sealed at M664 (position (1)) in all four subunits (Fig. 4F).

**Figure 4.**
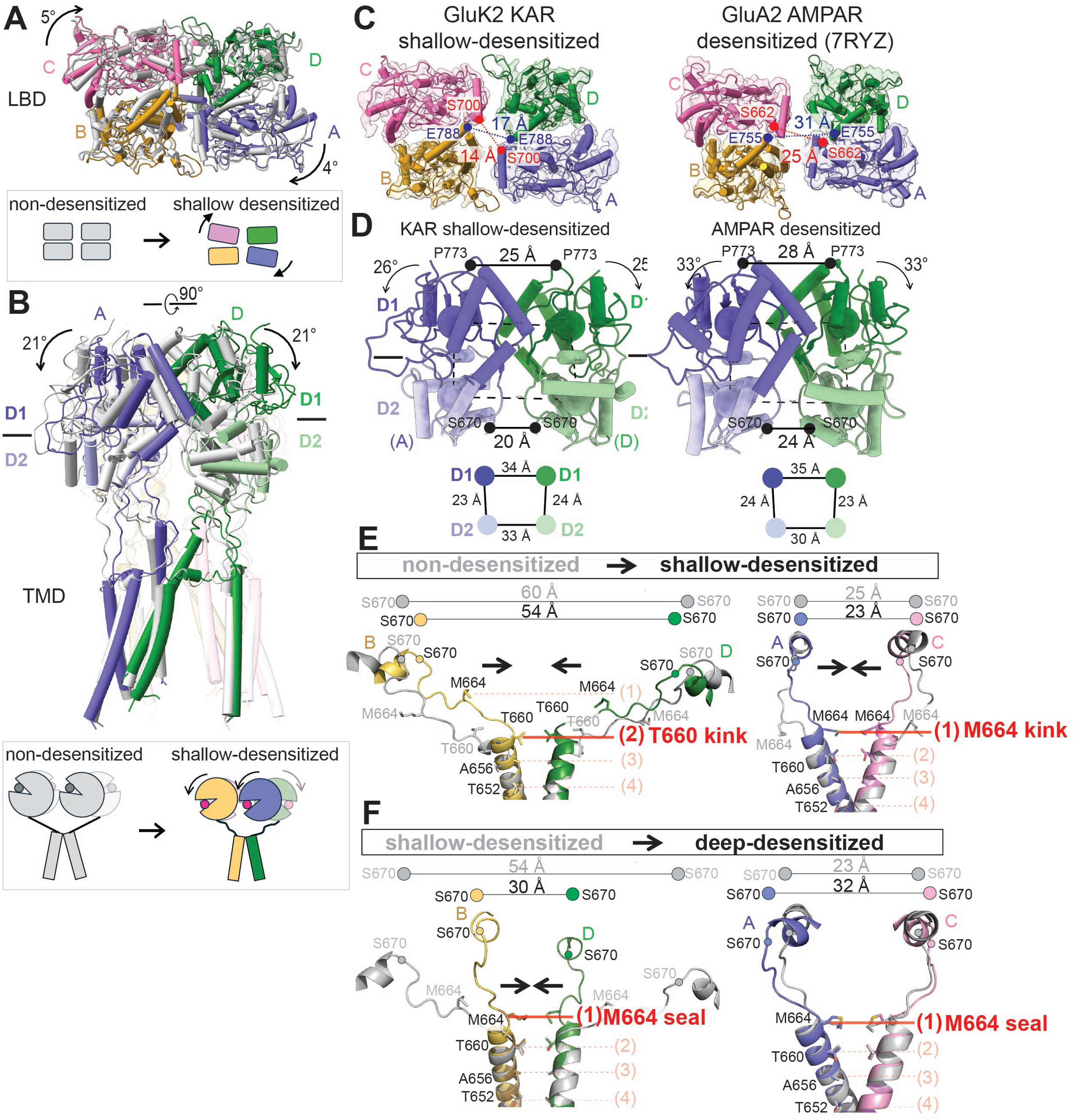
GluK2 stabilized in a shallow-desensitized-like conformations resembbles the desensitized conformation of AMPARs. **(A, B)** Structural comparison between the non-desensitized and shallow-desensitized conformations, viewed parallel (A) and perpendicular (B) to the membrane. Black arrows indicate the degree of LBD domain rotation. **(C, D)** Structural comparison of LBD layers in GluK2 K676C/N802C in the shallow-desensitized state and GluA2 AMPAR in the desensitized state (PDB code 7RYZ)^64^, viewed parallel (C) and perpendicular (D) to the membrane. Distances between S700 in the AC subunits and E788 in the BD subunits at the LBD dimer-dimer interface are shown in red and blue, respectively. The locations of the centers of mass (COM) of D1 and D2 are indicated by blue and green spheres, respectively, with the COM distances between D1 and D2 of subunit AD LBDs (shown as circles) displayed below. Arrows indicate D1 lobe rotations compared to the non-desensitized (KAR) or apo (AMPAR) states. Distances between the Cα atoms of P773 (D1-D1 distance) and S670 (D2-D2 distance) in the AD subunits are also indicated. **(E)** Comparison of LBD-TM3 linkers and TM3 helices in non-desensitized and shallow-desensitized conformations, showing the subunit BD pair (left) and subunit AC pair (right). S670 is shown as spheres in corresponding colors for the shallow-desensitized conformation and in gray for the non-desensitized conformation. The cross-dimer distances between the Cα atoms of S670 are indicated. Residues forming the ion channel pore are displayed as sticks. The locations of the TM3 gating hinge at T660 (subunit BD) and M664 (subunit AC) in the shallow-desensitized conformation are highlighted in red. The dotted line indicates the positions of pore-lining residues: (1) M664, (2) T660, (3) A656, and (4) T652. The shallow-desensitized conformation contains an additional half-turn of helices at T660 compared to the non-desensitized conformation. **(F)** Comparison of LBD-TM3 linkers and TM3 formation in shallow- and deep-desensitized GluK2, showing the subunit BD pair (left) and the subunit AC pair (right). S670 is represented as spheres in corresponding colors for the deep-desensitized state and in gray for the shallow-desensitized GluK2. In the deep-desensitized conformation, the ion channel is sealed at M664, completely closing the ion channel pore, in contrast to the shallow-desensitized conformation, which has a kink at T660 in the BD subunits.

### A large in-plane rotation of the LBD is required for proper ion channel closure during desensitization in GluK2 KAR

We next assessed the ion channel pore in these desensitized conformations. Although the extracellular domain in the shallow-desensitized GluK2 KAR and the desensitized GluA2 AMPAR adopt similar conformations, we observed distinct differences in their ion channel pore structures (Fig. 5A–C). The shallow-desensitized conformation has restriction points around T652, A656, and T660, which correspond to T617, A621, and T625 in the desensitized GluA2 (PDB code 7RYZ)^64^. Notably, the pore diameter at T660 and T652 in the shallow desensitized GluK2 KAR is wider compared to that in the desensitized GluA2 AMPAR. The ion channel pore in the transition state is also similar to that in the shallow-desensitized state, except for the presence of a narrower restriction point (<1 Å) around T652 (Fig. 5A). By contrast, all three deep-desensitized conformations exhibit multiple tight constrictions, particularly at the top of the gate region (Fig. 5A, D).

**Figure 5.**
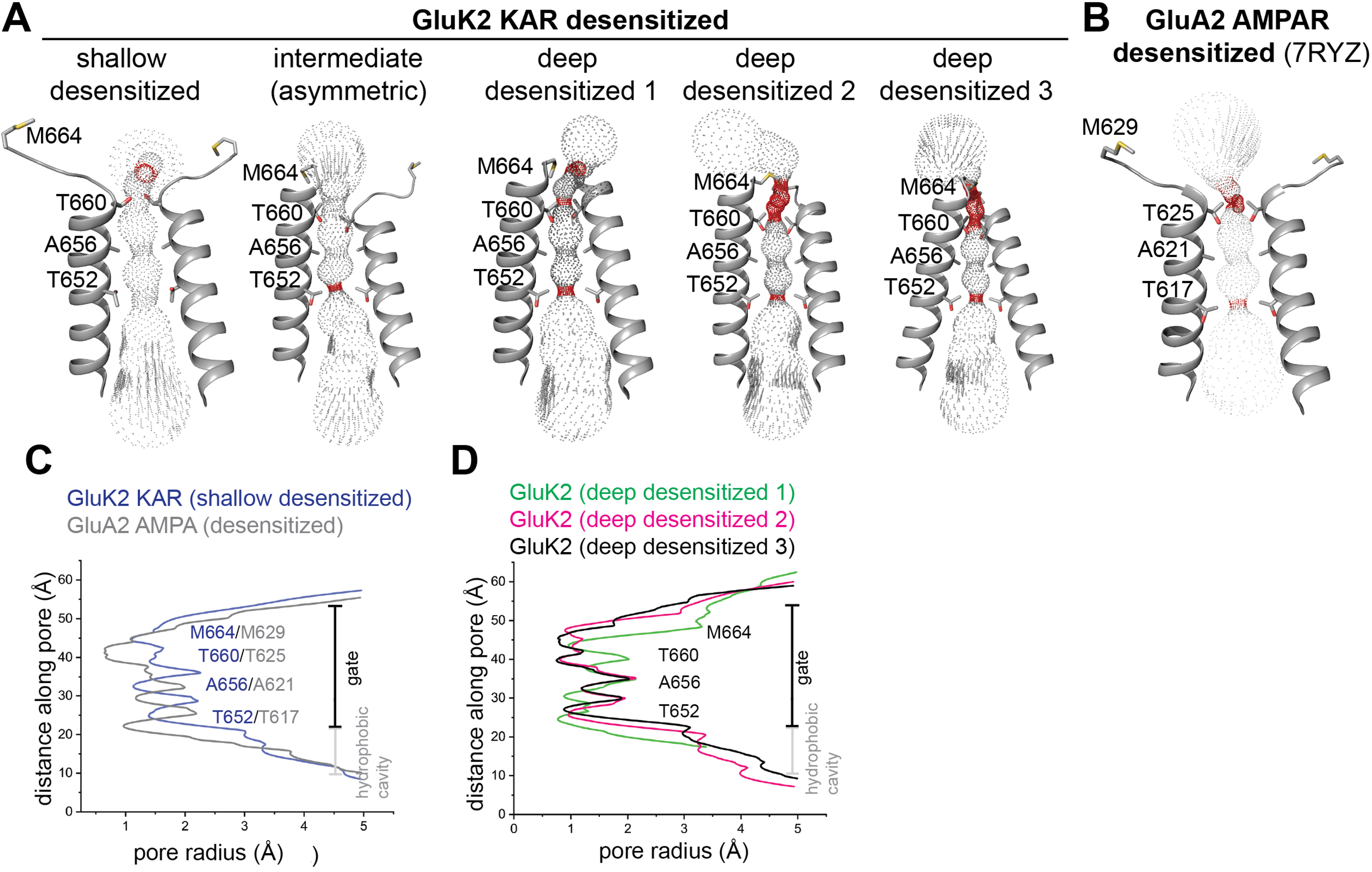
Ion channel pore of desensitized GluK2 KAR. **(A)** Permeation pathway dictating pore diameter of GluK2 K676C/N802C in desensitized states. Pore-delineating dots are colored according to pore radius: red for regions with a radius less than 1.1 Å and gray for regions with a radius greater than 1.1 Å. **(B)** Permeation pathway determining the pore diameter of the GluA2 AMPAR in a desensitized state (PDB code 7RYZ)^64^, using the same color code as (A). **(C–D)** Pore profile traces comparing GluK2 K676C/N802C KAR. Comparison between the shallow-desensitized conformation and the desensitized GluA2 AMPAR (PDB: 7RYZ) (C); and comparison among three deep-desensitized states (D).

As previously reported^28,29^, the ‘desensitization ring’ at the bottom of the LBD layer in these deep-desensitized conformations is stabilized by numerous polar interactions, supporting the four-fold symmetrical deep-desensitized conformations, whereas the shallow-desensitized and intermediate conformations exhibit disrupted ring formation, resulting in wider gate regions. Moreover, the large in-plane rotation of the LBDs in the BD subunits during desensitization twists the LBD-TM3 linker, creating an additional constriction at M664 at the top of the channel pore (Fig. 4F), which is not seen in AMPARs. We observe that the formation of the desensitization ring at the bottom of the LBD layer tightly closes the ion channel in the deep-desensitized conformations, whereas the shallow-desensitized state lacks this secure channel pore lock at T660 and M664 (Fig. 5A).

Based on these functional and structural observations, we hypothesized that the GluK2 KAR K676C/N802C mutant, which forms cysteine crosslinks, undergoes shallow rather than deep desensitization. This may allow the receptors to remain ion-permeable, exhibit a higher open probability, or recover quickly from desensitization, similar to AMPARs, thereby enabling more efficient activation of the receptors.

### K676C-N802C cysteine crosslinks partially blocks receptors from undergoing deep desensitization

To test our hypothesis, we first assessed the kinetics of the GluK2 K676C/N802C mutant using whole-cell patch-clamp electrophysiology. Remarkably, we observed steady-state currents of the cross-link mutant even at the highest concentration of glutamate (50 mM), which were not observed in the wild type GluK2 receptors (Fig. 6A). There were no significant differences neither in the current density at peak current between GluK2 WT and the mutant (GluK2 WT: 830 ± 50 pA/pF, n=6; GluK2 K676C/N802C: 700 ± 150 pA/pF, n=9), nor in the EC₅₀ for glutamate (GluK2 WT: 110 ± 30 µM, n=7; GluK2 K676C/N802C: 150 ± 50 µM, n=5), indicating that the cysteine crosslinks do not significantly affect agonist binding affinity or receptor activation (Fig. 6A-D). GluK2 K676C/N802C partially desensitized even by 0.01 mM glutamate, a concentration below the EC₅₀ (approximately 0.17 mM). This suggests that even partial occupancy is sufficient to initiate the conformational changes leading to desensitization, consistent with earlier findings^67,68^. While GluK2 WT desensitizes almost completely with a time constant of 8 ± 1 ms (n=14), the GluK2 K676C/N802C receptors in the presence of 0.01–50 mM glutamate exhibited approximately 60% desensitization, except when activated by 0.001 mM glutamate where only the steady-state current was observed with a decreased peak amplitude (Fig. 6E). Therefore, the degree of desensitization is not glutamate concentration dependent.

**Figure 6.**
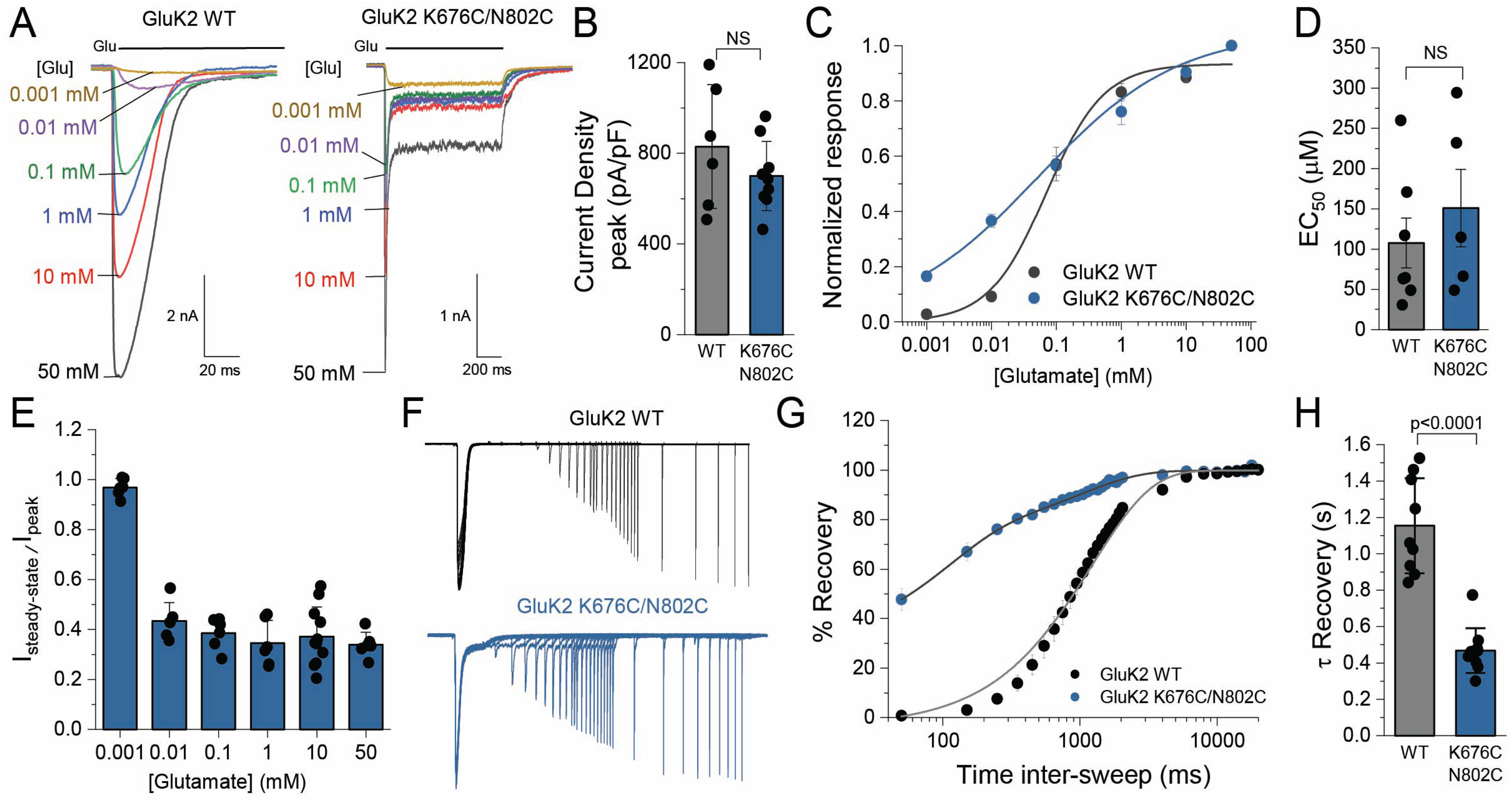
Functional characterization of GluK2 K676C/N802C receptor. **(A)** Representative whole-cell patch-clamp recordings of GluK2 WT (left) and GluK2 K676C/N802C (right) in response to varying concentrations of glutamate (0.001–50 mM). The black bar at the top of the recording represents the glutamate application. The cells were held at -70 mV. **(B)** Quantification of current density (pA/pF) at the peak current for GluK2 WT (gray) and K676C/N802C mutant (blue) activated by 10 mM glutamate. No significant differences were observed (NS, p > 0.05). **(C)** Glutamate dose-response curves for GluK2 WT (n=12) and K676C/N802C (n=8), normalized to the maximal response at 50 mM glutamate. Glutamate was applied for 1 s, and the peak current was measured at each concentration. All response were recorded at each concentrations using a single cell. **(D)** EC₅₀ values for glutamate activation of GluK2 WT (gray) and K676C/N802C (blue). No significant differences were detected (NS, p > 0.05). **(E)** Ratio of steady-state to peak current (I_steady-state_ / I_peak_) at various glutamate concentrations. The values of the steady-state current were taken at the end of the activation. **(F)** Representative current traces of GluK2 WT (black) and GluK2 K676C/N802C (blue) during a two-pulse protocol, showing the interval between glutamate exposures ranging from 50 ms to 20 s, highlighting altered desensitization kinetics in the mutant. **(G)** Recovery from desensitization at increasing inter-sweep intervals, fitted with one-term exponential functions for GluK2 WT (R²=0.99) and two-term exponential functions for GluK2 K676C/N802C (R²=0.99). GluK2 K676C/N802C exhibits a faster recovery than GluK2 WT. **(H)** Time constant (τ) of recovery from desensitization, showing a significant reduction in τ for the K676C/N802C mutant (p < 0.0001, two-sided two-sample t-test). The data for the WT was obtained from the fit in panel G, and for K676C/N802C, a mean τ was calculated using τ_mean_ = ((A1 τ1 + A2 τ2) / (A1 + A2)). Biological independent measurements: n=9 for GluK2 WT and n=9 for GluK2 K676C/N802C. Error bars represent Standard Deviation (SD). Statistical significance is indicated as NS (not significant) or with p-values where appropriate.

To assess the desensitization state of the crosslinked mutant, we next analyzed the rate of recovery from desensitization. As previously reported, recovery from desensitization in the GluK2 WT is extremely slow, requiring 1,150 ± 90 ms (n = 9) for full recovery. By contrast, the GluK2 K676C/N802C mutant exhibited a faster recovery from desensitization, characterized by two distinct components: τ₁ (100 ms, 60%) and τ₂ (1,000 ms, 40%) (n = 9). The τ mean of the mutant was 440 ± 40 ms, approximately three times faster than that of GluK2 WT (Fig. 6F–H). This shift brings its kinetics closer to those of AMPARs, which recover in the order of tens to hundreds of milliseconds (e.g. GluA1 and GluA2 wild-type recovery constants are approximately 200 ms and 9-50 ms, respectively)^69–71^. This result highlights the impact of cysteine cross-linking in accelerating desensitization recovery, potentially by stabilizing the dimer-of-dimers conformation.

### GluK2 KAR stabilized in a shallow-desensitized conformation remains ion-permeable

To assess whether the non-desensitized current and faster recovery kinetics result from crosslinking between the K676C and N802C, we performed whole-cell recordings in the presence of 5 mM dithiothreitol (DTT), applied for 7 minutes prior to the second glutamate activation. The peak intensity of both GluK2 WT and GluK2 K676C/N802C, in the presence and absence of DTT, showed no significant changes, confirming our earlier analysis that the crosslinks do not affect activation kinetics (Fig. 7A). By contrast, the application of 5 mM DTT for 7 minutes led to a >90% reduction in the steady-state current of GluK2 K676C/N802C (steady-state current prior to and after DTT application: 0.4 ± 0.1 nA and DTT: 0.03 ± 0.03 nA, respectively, n=8) (Fig. 7B). Additionally, recovery from desensitization of the DTT treated cells expressing GluK2 K676C/N802C was 1,040 ms ± 50 (n=5), similar to the WT (Fig. 7C, D). This experiment, together with the assessment of the single cysteine mutants, GluK2 K676C and GluK2 N802C (Fig. S2A), confirms that the observed non-desensitizing efficacy is specifically due to disulfide bond formation, which stabilizes the two-fold symmetrical non-desensitized or shallow-desensitized conformations as originally intended.

**Figure 7.**
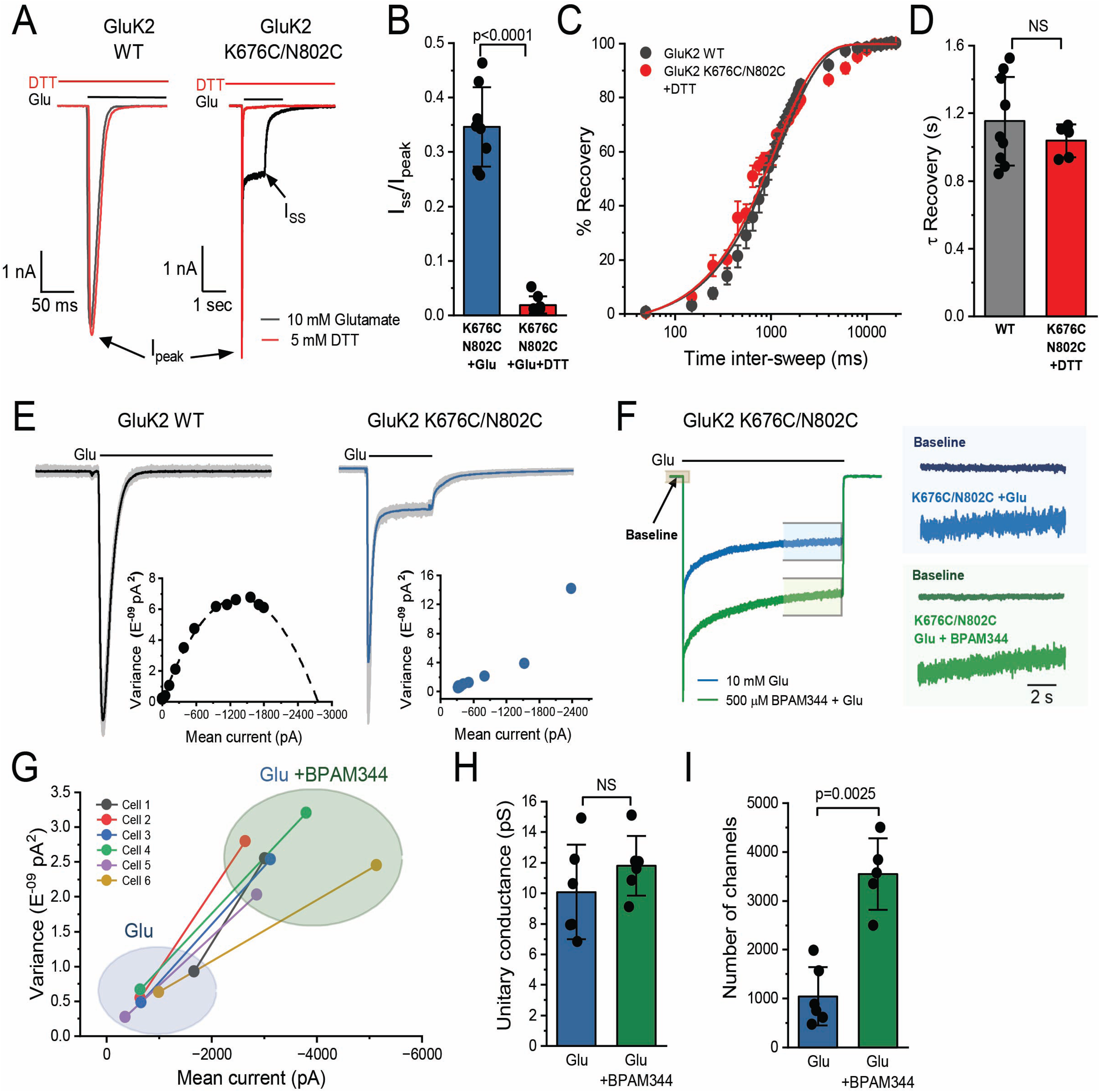
Cysteine crosslinking modulates the kinetics of GluK2 K676C/N802C. **(A)** Representative whole-cell patch-clamp recordings of GluK2 WT (left) and GluK2 K676C/N802C (right) in the presence of 10 mM glutamate before (black) and after treatment with 5 mM DTT for 7 minutes (red). **(B)** Quantification of the I_ss_/I_peak_ ratio in GluK2 K676C/N802C before (blue) and after (red) DTT treatment, showing a significant reduction in steady-state current post-treatment (p < 0.0001, two-sided two-sample t-test). **(C)** Recovery from desensitization at increasing inter-sweep intervals after treatment with 5 mM DTT. The GluK2 WT recovery curve from Figure 6, panel C is shown in black for comparison. The recovery kinetics of GluK2 K676C/N802C post-DTT treatment were fitted with a one-term exponential function (R² = 0.98), showing a recovery profile similar to WT. **(D)** Time constant (τ) of recovery from desensitization, showing that DTT treatment restores the recovery kinetics of GluK2 K676C/N802C to a level comparable to WT (NS, p > 0.05). **(E)** Representative whole-cell responses to 10 mM glutamate (200 ms, -70 mV; black bar) from HEK293T cells expressing GluK2 WT (left, average current in black, 60 individual responses in gray) and GluK2 K676C/N802C (right, average current in blue, 100 individual responses in gray). Inset: Current-variance relationship, with WT data fitting a parabolic function (dotted line), whereas the K676C/N802C mutant does not follow this trend. **(F)** Whole-cell recordings of GluK2 K676C/N802C in response to 10 mM glutamate in the absence (blue) and presence (green) of 500 µM BPAM344. Right panel: Representative noise traces at the end of glutamate application (∼10 s) under both conditions, indicated by blue and green boxes. Background noise (baseline) was taken from the current recording before glutamate application, as indicated by the gray box. **(G)** Variance-mean current relationship for individual cells in the presence of glutamate alone or with BPAM344, indicating an enhanced channel variance-current relationship upon BPAM344 application. **(H)** Unitary conductance determined by variance-mean analysis, showing no significant difference between two conditions with and without BPAM344 (NS, p > 0.05). (I) Estimated number of open channels, showing a significant increase upon BPAM344 application (p = 0.0025, two-sided paired t-test). Biologically independent measurements: n = 6. Data for glutamate and BPAM344 conditions were obtained from the same cell. Error bars represent standard deviation (SD). Statistical significance is indicated as NS (not significant) or with p-values where appropriate.

Next, we aimed to determine whether crosslinked receptors remain ion-conductive when exhibiting steady-state current. A key feature of KARs is their exceptionally small conductance and low open-channel probability, which makes it technically challenging to directly observe channel activity at the single-channel level, unlike other channels, including AMPARs^72,73^. Therefore, we conducted noise analysis to assess its conductivity. We first performed nonstationary fluctuation analysis (NSFA) on the desensitizing phase of macroscopic currents from GluK2 WT and GluK2 K676C/N802C, activated by glutamate pulses (10 mM, 100 ms). The variance of fluctuations around the expected current was analyzed. The weighted mean single-channel conductance of GluK2 WT was calculated as 23 ± 7.8 pS (n = 5), consistent with previous report^72^. While the current-variance relationship of the wild type was well-fitted by a parabolic function, the analysis of GluK2 K676C/N802C revealed an anomalous current-variance relationship, preventing the calculation of its conductance (Fig. 7E). However, the mutant generates a unique large fractional steady-state current. Thus, we next analyzed stationary fluctuation on the steady-state current of the desensitized receptors in the presence and absence of BPAM344.

To assess unitary conductance and the number of active channels, we analyzed the microscopic currents of GluK2 K676C/N802C activated by 10 mM glutamate for 30 seconds under desensitizing conditions (Fig. 7F). When activated by 10 mM glutamate, the mutant exhibited a unitary conductance of 10 pS—less than half of the wild-type conductance calculated by NSFA— and was similar to the lowest conductive level (O1) previously determined for KARs^72^. Variance-current analysis across individual cells showed that BPAM344 shifted receptor behavior toward higher variance and mean current values (Fig. 7G), suggesting an increase in the number of open receptors and their open probability. Interestingly, the conductance of the mutant activated by 10 mM glutamate was similar in the presence and absence of BPAM344 (GluK2 K676C/N802C, 10 mM glutamate: 10 ± 3 pS, n = 6; GluK2 K676C/N802C, 10 mM glutamate + 0.5 mM BPAM344: 10 ± 2 pS, n = 6) (Fig. 7H). Whereas the number of active channels in the presence of BPAM344 was approximately four-fold higher than in its absence (GluK2 K676C/N802C, 10 mM glutamate: 1,040 ± 600, n = 6; GluK2 K676C/N802C, 10 mM glutamate + 0.5 mM BPAM344: 4,080 ± 460, n = 6) (Fig. 7I). In summary, these experiments indicate that desensitized GluK2 K676C/N802C KARs remain conductive at the single channel level, although with significantly lower conductance compared to wild-type channels activated by glutamate.

## Discussion

Receptor desensitization is a fundamental property of ligand-gated ion channels which depends on agonist, receptor subtypes, and auxiliary proteins. Recent advances in iGluR research have revealed significant differences in the kinetics and conformational dynamics of KARs during desensitization compared to other iGluRs^1,25,28,29,31,33^. The desensitized conformation of KARs is characterized by the quasi-four-fold symmetrical LBDs, by contrast to typical desensitized iGluRs, which remain in two-fold symmetrical conformations. In addition, functionally, the majority of KARs enter deep-desensitized states, with recovery from desensitization occurring 5- to 130-fold slower than AMPARs. By contrast, most AMPARs typically adopt shallow-desensitized states, exhibiting deep desensitization only under certain conditions^74,75^. In this study, we employed a classical cysteine crosslinking approach to answer the fundamental question why KARs require such dramatical conformational changes for proper channel closure and complete desensitization, which are not conserved in other iGluRs.

In the previously determined structure of BPAM344- and ConA-bound GluK2 in a highly ion-conductive open state, ConA binds both the ATD and LBD, occupying the space between these two layers^27^. As a result, the organization of the ATD-LBD layers upon activation remains unclear. In this study, we analyzed the extracellular domain organization in the absence of ConA. The assembly of the extracellular domains of our glutamate- and BPAM344-bound GluK2 K676C/N802C in the non-desensitized state resembles that of GluK2 in its open state. Interestingly, in the absence of ConA, the ATD-LBD distance in the non-desensitized state of GluK2 K676C/N802C is wider. Thus, upon activation, the ATD layer is elevated and undergoes a counterclockwise in-plane rotation. By contrast, the apo and desensitized receptors adopt more compact extracellular domain conformations, with ATD and LBD layers positioned closer. Since ATD-deleted KARs remain functional as ion channels^27,76^, further investigation is needed to determine how these conformational changes in the ATD layer contribute to receptor activation. In addition, we also observed the TM3, which form the gating region of the ion channel pore, kinks at the position between the kinking observed in the apo and the open structure (Fig. 8), along with rotational movement of TM3. This observation suggests that TM3 kinking at a higher position (A656/E662) within the gating region is insufficient for channel opening—particularly full opening with high conductivity—which instead requires TM3 rotation and kinking at a lower position (L655) in all four subunits (Fig. 8).

**Figure 8.**
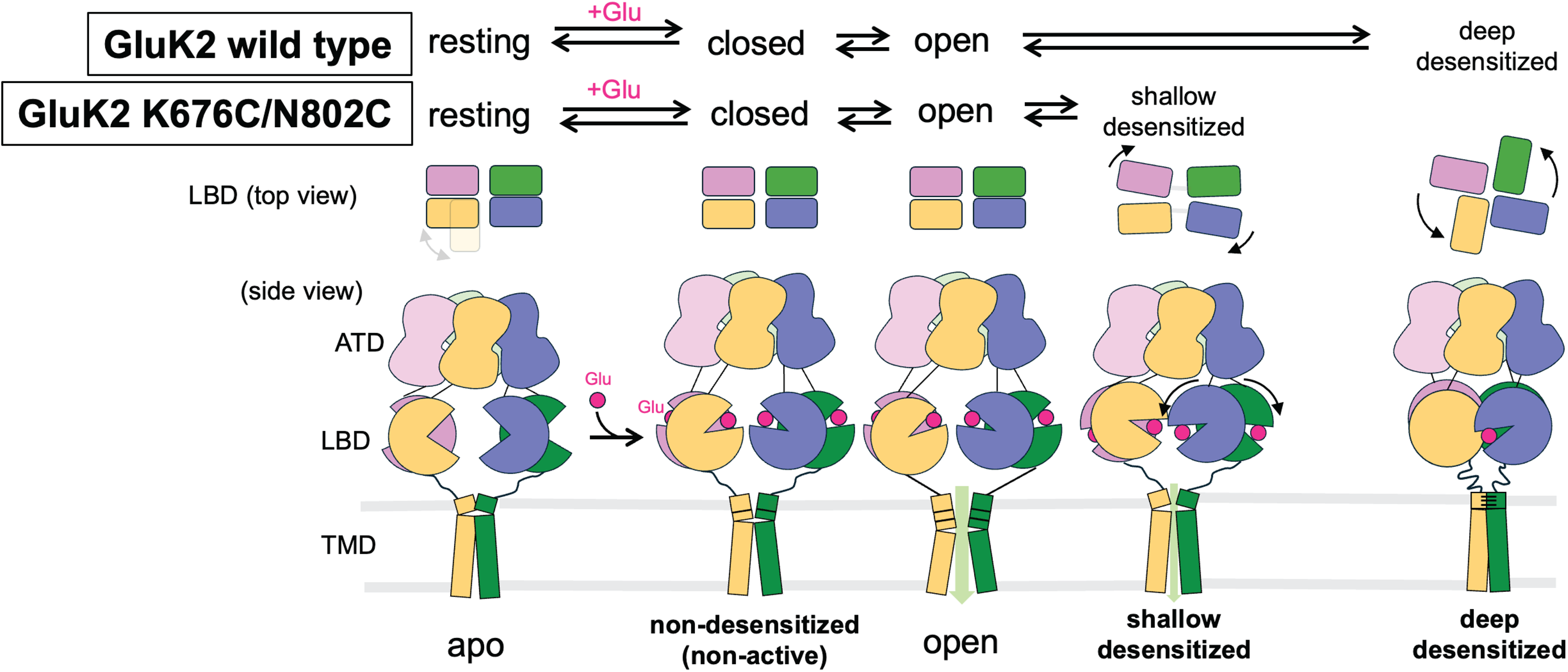
Schematic representation of conformational rearrangements during activation and desensitization. The proposed model illustrates how conformational changes in the LBD layers during desensitization regulate ion channel permeation. Wild-type GluK2 typically transitions into a deeply desensitized state, adopting a stabilized, classically four-fold symmetrical conformation. By contrast, receptors that stabilize in a two-fold symmetrical, desensitized AMPAR-like conformation, maintained by engineered disulfide bonds, remain conductive, although their conductivity is lower than that of receptors in the open state. The position of the gating kink differs among the non-desensitized, open, and shallow-desensitized conformations. Additionally, the ATD layers elevate and rotate in both the non-desensitized (non-active) and open-state GluK2 conformations.

Conversely, the glutamate-bound GluK2 K676C/N802C structures represent shallow- and deep-desensitized conformations as well as an intermediate state between these two desensitized states. The shallow-desensitized conformation demonstrates that two inter-dimer crosslinks stabilize receptors in a two-fold symmetrical conformation with D1-D1 ruptured LBDs, similar to the desensitized state of AMPARs^42,44,63–65^. While most previously determined AMPAR structures in desensitized states are stabilized in the classical two-fold symmetrical conformation, the partial agonist, fluorowilliardiine-bound GluA2 AMPAR, in a desensitized state exhibited an asymmetrical conformation with the LBD of subunit D undergoing a large conformational rearrangement similar to our GluK2 in the intermediate-state conformation^77^. Additionally, recent studies have shown that extracellular protons influence extracellular domains and alter LBD conformation, causing minor in-plane rotation of LBDs within the submit B and D, although these changes are not as pronounced as those observed in the deep-desensitized KAR conformations^70,71,78^. Nevertheless, adopting the four-fold symmetrical deep-desensitized conformation is not energetically favorable for AMPARs during desensitization^74^, whereas KARs exhibit greater stability in this four-fold conformation. Previous mutagenesis studies demonstrated that various mutations in the D2 lobe of KARs alter desensitization and recovery from desensitization, but no mutation completely blocked desensitization unlike AMPARs^62^. This suggests that multiple non-conserved residues contribute to the stability of the four-fold desensitized conformation of KARs, despite the overall architectural similarity between AMPAR and KAR LBDs. Our electrophysiological experiments demonstrate that cysteine crosslinking prevents the receptors from undergoing complete desensitization, resulting in significantly stronger and faster recovery—resembling the kinetics of AMPARs rather than those of GluK2 WT. Upon treatment with 5 mM DTT, which breaks the disulfide bonds in the GluK2 K676C/N802C mutant, recovery from desensitization resembled that of the WT, indicating a correlation between LBD conformation and desensitization recovery in GluK2 KAR.

Furthermore, we aimed to understand why KARs have to form the four-fold symmetrical arrangement upon desensitization. Previous studies have shown that in the “desensitized ring”, a network of polar interactions stabilizes the deep-desensitized conformation of the LBD layer. This, in turn, helps to stabilize the conformation of the TMD layer, thereby reinforcing receptor desensitization and influencing the kinetics of desensitization recovery^28,29^. In the shallow-desensitized conformation, lacking desensitization ring formation with the two-fold symmetric LBDs, we observed that the ion-channel is dilated by contrast to channels observed in desensitized AMPARs and fully/deep desensitized KARs, which show much narrower channel pores throughout the gating region with the channel pore tightly sealed on the top of the gating region. Previously, a conducting desensitized state of AMPAR and acetylcholine receptors has been reported^41,79^. In addition, it was microscopically shown that the partially desensitized KARs are conductive, as defined by the number of desensitized subunits per KAR tetramer^80^. The structural observation together with the characteristic steady-state current of the GluK2 K676C/N802C indicates that the receptors in the shallow-desensitized state could form weakly conductive channels. Indeed, our noise analysis demonstrates that the desensitized GluK2 K676C/N802C receptors, exhibiting steady-state current, remains conductive at the single-channel level; however, its conductance is approximately 30% weaker than the GluK2 WT, regardless of the presence of the positive allosteric modulator BPAM344. In addition, the crosslinked receptors exhibit up to ten-fold faster recovery from desensitization, whereas reducing the disulfide bonds with DTT restores currents similar to those of the GluK2 WT.

Taken together, this study answers the fundamental question by demonstrating that KARs absolutely require large conformational rearrangements of the LBD layer and stabilization in a four-fold symmetrical conformation to achieve proper channel closure and full desensitization into a non-conductive state (Fig. 8). This structural feature distinguishes KARs from other iGluR subfamilies, in which such symmetry and conformational changes are not observed.

## Materials and Methods

### Plasmid construction

For Cryo-EM studies, the construct used for the structural studies was full-length rat GluK2 KAR (GenBank 54257, Uniprot code P42260) with two mutations of N802C and K656C. The gene was cloned into the pFW vector (gift from Dr. Hiro Furukawa at Cold Spring Harbor Laboratory) and fused to a C-terminal human rhinovirus 3C protease recognition site and a 1D4 affinity tag. The construct was referred to as GluK2 N802C/K656C.

### Cell culture

For structural studies, HEK293S GnTI^-^ cells were grown to a density of 3.2 × 10^6^ cells / ml in FreeStyle 293 medium at 37°C and 8% CO_2_ supplemented with 2 % fetal bovine serum. The rGluK2 N802C/K676C mutant bacmid and baculovirus were generated as previously described. The P1 and P2 viruses were produced in Sf9 cells. Cells were infected with the baculovirus harboring GluK2 EM and incubated at 37°C for 12 hrs. Cells were supplemented with 10mM sodium butyrate and 20 µM DNQX and temperature was shifted to 30°C at 12hrs post-infection. The cell culture was incubated additional 60 hrs and harvested by low-speed centrifugation at 5,000 g for 20min and stored at -80°C until use.

For electrophysiology experiments, human embryonic kidney 293T (HEK293T) cells were cultured in Dulbecco’s modified Eagle medium (DMEM, Corning) supplemented with 10% FBS at 37°C in a 95% O₂–5% CO₂ atmosphere. Wild-type rat GluK2 and rat GluK2 K676C/N802C DNAs were cloned into the pCAG-IRES-EGFP vector (Addgene plasmid #119739). HEK293T cells were transfected with 1 µg/µl cDNA using the TransIT2020 transfection reagent (Mirus) according to the manufacturer’s instructions. Cells expressing GluK2 wild-type were transfected for 24 hours, while those expressing K676C/N802C were transfected for 18 hours. After transfection, cells were dissociated using Accutase (Innovative Cell Technologies, Inc.), resuspended, and plated onto 35 mm poly-D-lysine-coated dishes (Neuvitro). Electrophysiological recordings were performed 2–4 hours later^81^.

### Electrophysiological recordings

All whole-cell recordings were performed using HEKA EPC10 amplifiers (HEKA Elektronik, Lambrecht, Germany) with thin-wall borosilicate glass pipettes (2–5 MΩ) coated with dental wax to reduce electrical noise. Currents were recorded at a holding potential of −70 mV, with a sampling frequency of 10 kHz, and filtered at 2.6 kHz. The external solution contained (in mM): 145 NaCl, 2.5 KCl, 1.8 CaCl₂, 1 MgCl₂, 5 glucose, and 5 HEPES. The internal pipette solution contained (in mM): 105 NaCl, 20 NaF, 5 Na₄BAPTA, 0.5 CaCl₂, 10 Na₂ATP, and 5 HEPES. The pH and osmotic pressure of the external and internal solutions were adjusted to 7.4 and 300–290 mOsm/kg, respectively. L-glutamate was applied using theta glass tubing mounted on a piezoelectric stack (MXPZT-300 series, Siskiyou) driven by a HEKA EPC10 amplifier. Typical 10–90% rise times were 250–300 µs, measured from junction potentials at the open tip of the patch pipette after recordings. Data acquisition was performed using PULSE software (HEKA Elektronik, Lambrecht, Germany).

### Data analysis

The macroscopic rate of desensitization (τ_desensitization) was measured by fitting an exponential function to the decay of current from ∼80% of its peak amplitude (I_peak) to baseline in recordings where glutamate was applied for 1 second. For the double mutant N802C/K676C, only the first exponential component, which exhibited desensitization kinetics, was considered, while the steady-state current was omitted. Desensitization kinetics were fitted using a single-exponential, one-term Levenberg-Marquardt fitting approach.

To evaluate recovery from desensitization, two protocols were employed: (1) a two-pulse application of 10 mM glutamate for 100 ms per pulse, with interpulse intervals increasing in 100 ms steps from 50 ms to 2 s; and (2) a second protocol with interpulse intervals increasing in 2 s steps, ranging from 50 ms to 20 s. In all cases, the first peak was considered the control (100% activation), while the second peak represented recovery at a given time point. Data from both protocols were combined, and a recovery curve was generated, plotting the percentage of recovery against time. The experimental data were fitted to one-term exponential model, from which the recovery time constant (τ) was derived.

To analyze the EC₅₀ for glutamate in wild-type (WT) and mutant receptors, varying concentrations of L-glutamate were applied to a single cell using a theta glass tubing mounted on a piezoelectric stack (MXPZT-300 series, Siskiyou), driven by a HEKA EPC10 amplifier, for 1 second. One input of the theta glass tubing was connected to the control solution, while the other was connected to a six-inlet manifold for different L-glutamate concentrations. A 2-minute wash was used between concentration changes to ensure complete exchange before applying the next solution. The L-glutamate concentrations tested were 50, 10, 1, 0.1, 0.01, and 0.001 mM, with 50 mM serving as the normalization reference. The normalized data are presented as mean ± S.E.M. in a dose-response curve, with response on the y-axis and concentration on the x-axis. Experimental data from independent biological measurements (each defined as data from one cell with at least five concentration responses) were fitted to the Hill equation. The resulting EC_50_ values were then averaged and presented as mean ± S.E.M.

### Non-stationary fluctuation analysis (NSFA)

NSFA was performed on the decay phase of currents evoked by 200 ms applications of 10 mM glutamate (60–150 successive applications). Currents were sampled at 50 kHz and filtered at 5 kHz. The variance and mean current were calculated for each set of currents and then binned into 15 bins. The single-channel current and total number of channels (N) were determined by plotting the binned variance against the binned mean current and fitting the data with a parabolic function:

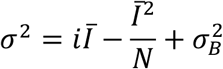

Where 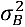 represents the background variance, which was calculated from the baseline before glutamate application. The 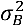 value was used to correct the variance, while the mean I_B_ was used to correct the mean current.

### Stationary fluctuation analysis (SFA)

HEK293T cells transfected with GluK2 K676C/N802C were activated with 10 mM glutamate for 30 seconds, followed by a 2-minute incubation with 500 µM BPAM344. The cells were then reactivated with 10 mM glutamate + 500 µM BPAM344. Currents were sampled at 10 kHz and filtered at 2 kHz. For each condition (glutamate alone and glutamate + BPAM344), the last 10 seconds of the steady-state current were used to calculate the variance and mean current. Since the stationary current represented approximately 35% of the peak current, we assumed that the open probability (P₀) in the steady-state current was P₀ < 1. The unitary current (i) was then estimated using the following equation:

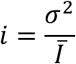

The unitary current was then used to estimate the unitary conductance (γ) as follows:

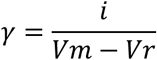

where Vₘ is the holding potential (−70 mV) and the reversal potential (Vᵣ) is 0 mV.

The number of channels (N) was estimated using the equation:

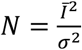

Statistical analyses were performed using OriginLab 2025. Statistical significance was calculated using a two-sided two-sample t-test, with significance assumed if P < 0.05.

### Protein expression and purification

Cell pellets were resuspended in ice-cold lysis buffer containing 20 mM Tris-Cl (pH 8.0), 150 mM NaCl, 0.2 mM PMSF, 20 µg/mL DNase I, and 20 µM DNQX. Cells were lysed by sonication using a Qsonica sonicator with an amplitude of 40% for 10 minutes total (10 s on, 10 s off). The lysate was centrifuged at 7,000 g for 20 min to remove cell debris and unbroken cells, and the supernatant was subjected to ultracentrifugation at 200,000 g for 30 min to pellet the cell membranes. The membrane pellet was homogenized and solubilized by gentle nutation for 2 hours at 4 °C in buffer consisting of 20 mM Tris-Cl (pH 8.0), 150 mM NaCl, 1% Dodecyl-β-D-Maltosid (w/v, DDM), and 20 µM DNQX, followed by centrifugation at 200,000 g for 30 min to remove insoluble material. The supernatant was incubated with Rho 1D4 affinity resin overnight at 4 °C. The protein-bound resin was extensively washed with 100 mL of buffer consisting of 20 mM Tris-Cl (pH 8.0), 500 mM NaCl, 1 mM DNQX, and 0.02% DDM at room temperature, and then slowly washed with 200 mL of buffer consisting of 20 mM Tris-Cl (pH 8.0), 300 mM NaCl, and 0.02% DDM at room temperature at a rate of approximately 0.3 ml/min. The protein was eluted in 50 mL of buffer with 20 mM Tris-Cl (pH 8.0), 300 mM NaCl, 0.02% DDM, and 0.25 mM 1D4 peptide. The eluted protein was concentrated and injected into a Superose 6 10/300 GL size-exclusion column equilibrated with buffer containing 20 mM Tris-Cl (pH 8.0), 150 mM NaCl, and 0.02% DDM. The peak fractions corresponding to tetrameric GluK2 were pooled and concentrated to 3.4 mg/mL for grid freezing.

### Grid preparation and cryo-EM data

The cryo-EM grids were prepared as follows. UltrAuFoil R1.2/1.3 300 mesh gold grids were glow-discharged for 50 seconds before sample application. A final concentration of 1 mM glutamate was added to the sample, and then a droplet of 3 µL of protein sample was rapidly applied onto the grids. For the BPAM344/glutamate sample, proteins were pre-incubated with 500 µM BPAM344 for 30 minutes on ice, followed by the addition of 1 mM glutamate to the sample. 3.5 µl of the protein sample was applied to the grids, and the grids are frozen within 10 seconds. An FEI Vitrobot Mark IV (Thermo Fisher Scientific) was used to plunge-freeze the grids into liquid ethane after sample application at 4°C and 100% humidity, with a blot time of 3 seconds and a blot force of −5.

All cryo-EM data were collected on a Titan Krios 300 kV microscope (Thermo Fisher Scientific) equipped with a GIF-Quantum energy filter (Gatan) with the slit set to 20 eV and a Gatan K3 Summit direct electron detection camera (Gatan) at Case Western Reserve University. Raw movies were collected using EPU in super-resolution mode, with a physical image pixel size of 1.06 Å and a defocus range of −0.8 to −1.6 µm. The total dose of around 50 e⁻/Å² was used for each movie, distributed across 50 frames for a total exposure time of approximately 3.2 seconds.

### Cryo-EM image processing

For dataset **a** (GluK2 K676C/N802C supplemented with 1 mM glutamate and 500 µM BPAM344), the same motion correction procedure was applied using Patch Motion Correction in CryoSPARC4. CTF calculations were also carried out on the motion-corrected images using Patch CTF Estimation in CryoSPARC4. Micrographs unsuitable for further analysis were manually removed. Initial particle selection from 1000 micrographs was done with Blob Picker, followed by 2D classification to generate good templates. Template Picker was used to pick 903,654 particles, which were extracted with a pixel size of 2.12 Å. After several rounds of 2D classification, 612,540 particles were selected for initial 3D model generation via ab initio reconstruction. A total of 352,269 particles were selected and re-extracted with a pixel size of 1.06 Å following multiple rounds of heterogeneous refinement. NU-refinement for each class resulted in maps with resolutions of 6.16 Å, 5.80 Å, 4.14 Å, 4.00 Å, 4.00 Å, and 3.83 Å, respectively.

For dataset **b** (GluK2 K676C/N802C supplemented with 1 mM glutamate), the raw movie stacks were motion-corrected using Patch Motion Correction in CryoSPARC4. The Contrast Transfer Function (CTF) was calculated from the motion-corrected images using the Patch CTF Estimation in CryoSPARC4. Micrographs deemed unsuitable for further analysis were removed through manual inspection. Initial particle selection was performed using Blob Picker from 2,000 micrographs, followed by 2D classification to generate good templates. Subsequently, 1,005,623 particles were picked using Template Picker and extracted with a pixel size of 2.12 Å. After several rounds of 2D classification, 765,893 particles were selected to generate the initial 3D model through ab initio reconstruction. A total of 277,369 particles were selected and re-extracted with a pixel size of 1.06 Å after multiple rounds of heterogeneous refinement. NU-refinement for each class yielded maps with resolutions of 3.92 Å, 4.35 Å, 3.85 Å, and 3.82 Å, respectively. To improve the structure resolution for classes present in both datasets, particles from each class were combined and subjected to several rounds of heterogeneous refinement to remove junk particles. Separate NU-refinements for each class generated maps with resolutions of 3.67 Å, 4.02 Å, 3.81 Å, and 3.74 Å, respectively. For Class 2, focused masks around the ATD and LBD-TMD regions were created separately and used for local refinements of the corresponding regions. As a result, 3.42 Å reconstruction of the ATD region and 3.86 Å reconstruction of the LBD-TMD region were obtained through local refinement with focused masks. For the non-desensitized state, particle subtraction was performed for both ATD-focused and LBD-TMD-focused heterogeneous refinement. After several rounds of heterogeneous refinement, 80,325 particles were used for subsequent NU-refinement, resulting in a 3.70 Å resolution of the ATD 3D reconstruction. A total of 117,714 particles were subjected to NU-refinement for the LBD-TMD region, generating 3.93 Å maps. The Gold-standard Fourier Shell Correlation (FSC) 0.143 criteria were used to estimate the map resolution for each class. For the shallow-desensitized state, focused masks around the ATD and LBD-TMD regions were created separately and used for local refinements of the corresponding regions. As a result, 3.42 Å reconstruction of the ATD region and 3.86 Å reconstruction of the LBD-TMD region were obtained through local refinement with focused masks.

### Model building and refinement

The model building for the non-desensitized class was initiated by individually fitting rigid-body components of the LBD extracted from the BPAM344-, and ConA-bound GluK2 structure (PDB code 9B36), and the TMD and ATD from the BPAM344 bound GluK2 structure (PDB code 8FWQ) into the cryo-EM map using Chimera. Subsequent manual adjustments, including fitting the backbone and side chains of the ATD-LBD and LBD-TM3 linkers into the density, as well as deleting unresolved side chains and making subtle local adjustments to minimize clashes, were performed in Coot.

For the shallow-desensitized class, the single LBD, entire ATD, and TMD were separately extracted from the BPAM344 bound GluK2 structure (PDB code 8FWQ) and manually docked into the corresponding cryo-EM map using Chimera. Further manual adjustments of the backbone and side chains were completed in Coot.

For intermediate and deep-desensitized classes, the single LBD, entire ATD, and TMD were extracted from the deep-desensitized GluK2 structure (PDB: 5KUF) and individually docked into the corresponding cryo-EM maps for each class using Chimera. Additional manual adjustments to the backbone and side chains were completed in Coot.

All four initial models were then refined in real space using Phenix against the corresponding cryo-EM maps. The models were validated through comprehensive validation procedures in Phenix. Glutamate ligands and BPAM344 compounds were extracted from the cryo-EM structure (PDB code 9B36) and manually fitted into the corresponding densities in Coot.

### Key source table

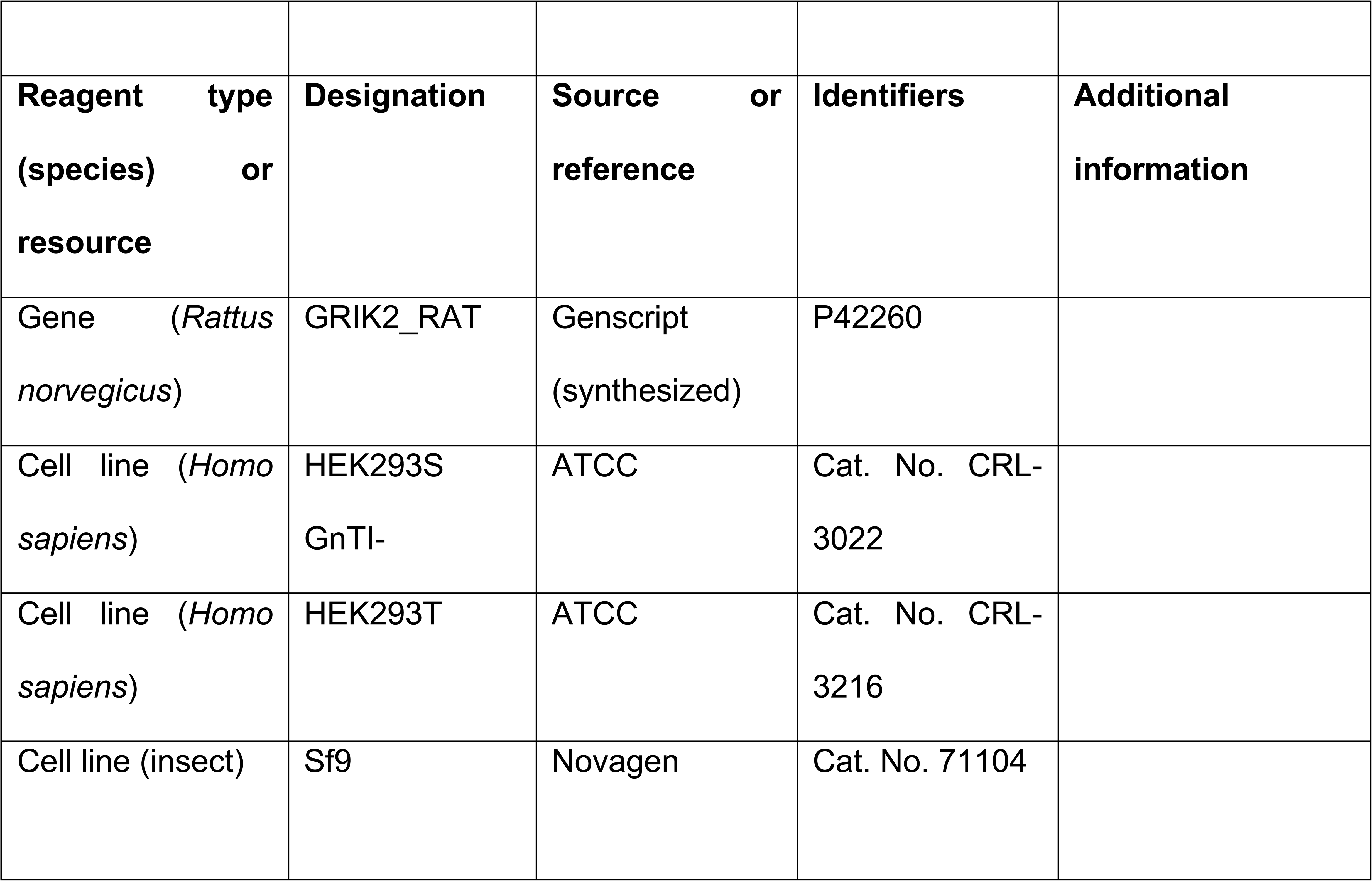
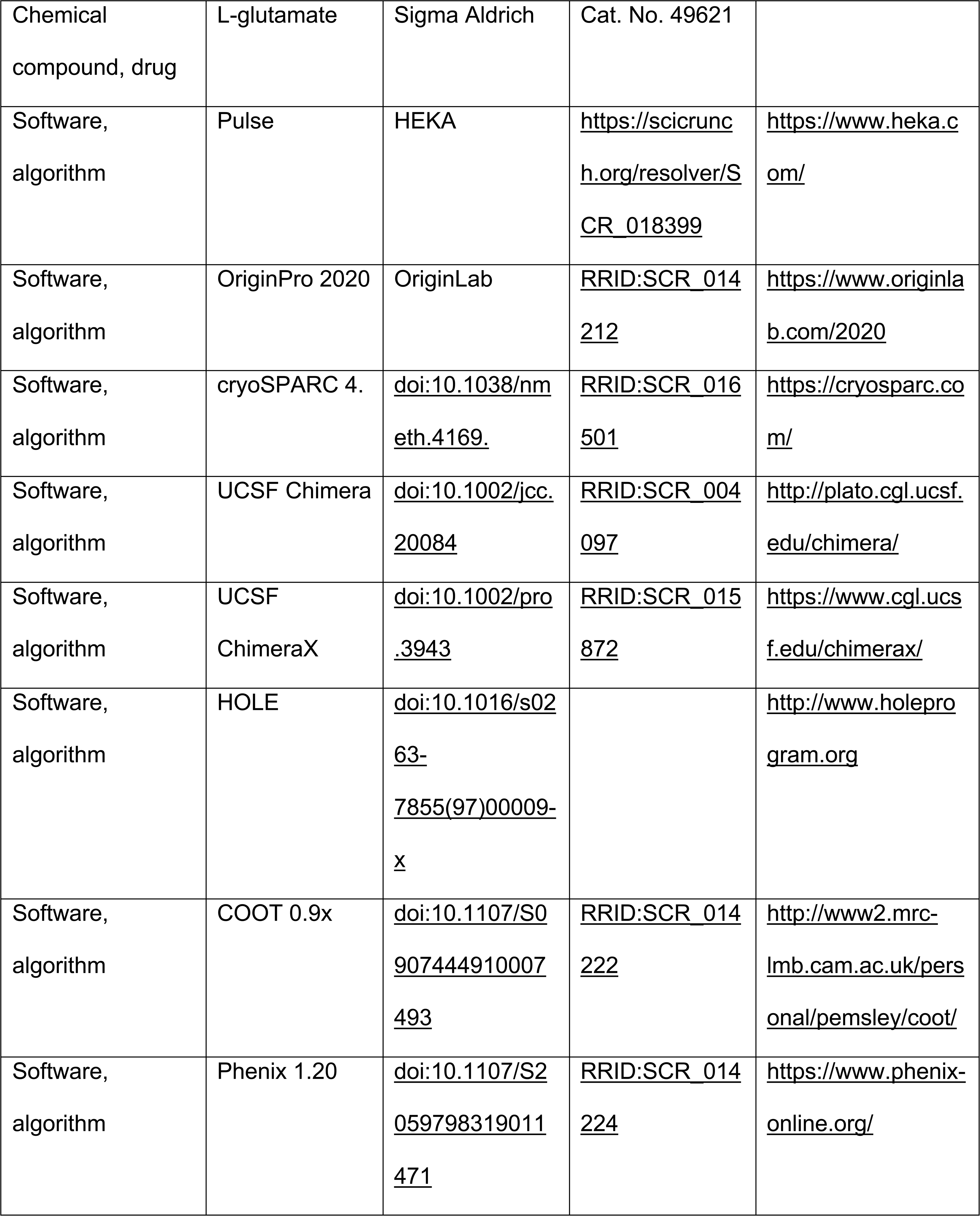
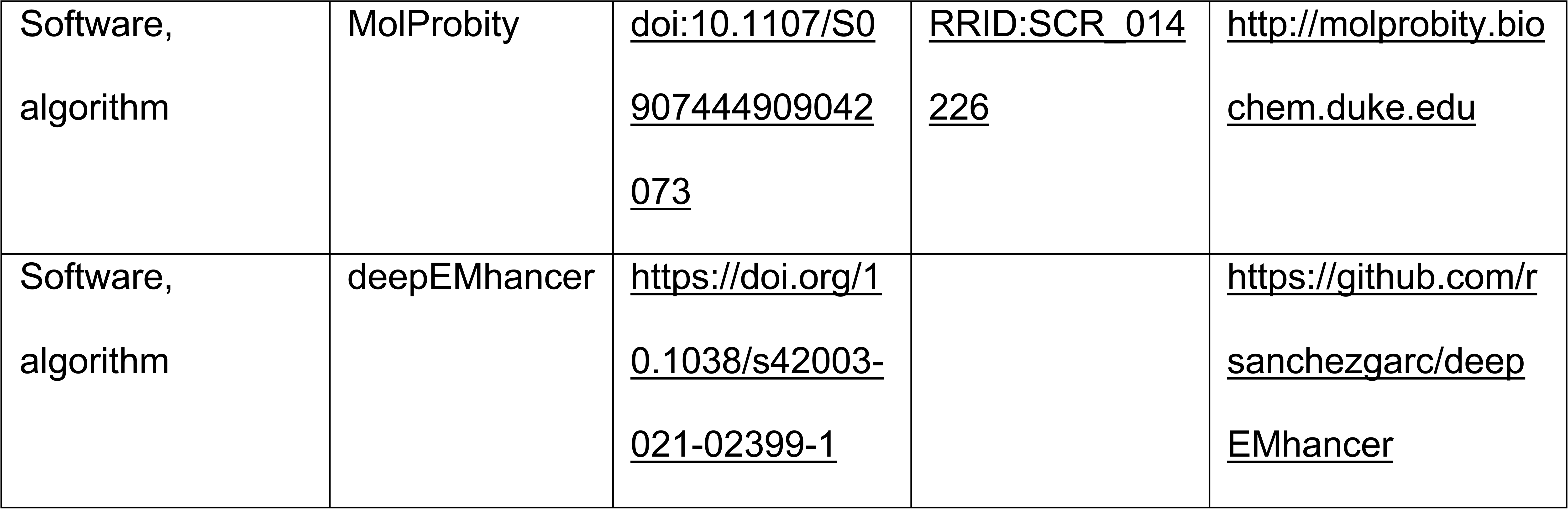

## Acknowledgment

We thank Dr. Matthias Buck (CWRU) for discussion and critical feedback on the manuscript. We thank Drs. Kunpeng Li and Kyle Whiddon at Case Western Reserve University (CWRU) for their cryo-EM technical support, and the High-Performance Computing Resource CWRU Core facility for their computational support. We are grateful to Drs. Corey Smith and Shyue-An Chan at CWRU for their technical assistance with patch-clamp electrophysiology. We thank Ashlee Hoffman (CWRU) for proofreading the manuscript. This work was funded by the National Institute of Health (1R35GM147266-01 to N.T.), Whitehall Foundation (N.T.), and American Heart Association https://doi.org/10.58275/AHA.23POST1019193.pc.gr.174253 (S.G.).

## Author contributions

C.Z., S.G., and N.T. conceived the project and designed the experimental procedures. C.Z. designed the constructs, conduced the protein expression and purification, and performed the EM data collection and data analysis. S.G. conducted electrophysiological recordings. C.Z., S.G., and N.T. wrote the manuscript. All authors reviewed the final draft.

## Competing interests

The authors declare no competing interests.

## Data and materials availability

Cryo-EM density maps and atomic coordinates for GluK2-apo and GluK2-domoate were deposited in the electron microscopy data bank under the accession codes listed below.

| Name | Accession code |
| --- | --- |
| GluK2 ATD in the non-desensitized state | PDB 9MZQ, EMD-48765 |
| GluK2 LBD-TMD bound to BPAM344 and glutamate in the non-desensitized state | PDB 9MZR, EMD-48766 |
| GluK2 bound to BPAM344 and glutamate in the non-desensitized state, composite map | PDB 9MZS, EMD-48767 |
| GluK2 ATD in the shallow-desensitized state | PDB 9MZN, EMD-48762 |
| GluK2 LBD-TMD in the shallow-desensitized state | PDB 9MZO, EMD-48763 |
| GluK2 in the shallow-desensitized state, consensus map | PDB 9MZM, EMD-48761 |
| GluK2 in the shallow-desensitized state, composite map | PDB 9MZP, EMD-48764 |
| GluK2 in the intermediate state | PDB 9MZL, EMD-48760 |
| GluK2 in the deep-desensitized state 1 | PDB 9MZK, EMD-48759 |
| GluK2 in the deep-desensitized state 2 | PDB 9MZJ, EMD-48758 |
| GluK2 in the deep-desensitized state 3 | PDB 9MZI, EMD-48757 |

---

**Figure S1.** Design of the double cysteine crosslink mutant of GluK2. **(A)** Schematic illustration of the KAR subunit topology. Each KAR subunit is composed of four domains: ATD, LBD,TMD, and CTD. The LBD contains two segments termed D1 (upper lobe) and D2 (lower lobe). Agonists bind to the interface between the D1 and D2 lobes of the LBD. **(B)** Homology model of the active GluK2 KAR, generated based on the active GluA2 AMPAR (PDB code 5WEO) as a template. **(C)** The inter-protomer interface between subunit A and subunit B in the homology model. **(D)** The distance between residues K676 and N802 in the BPAM344-, concanavalin A-, and glutamate-bound open/active GluK2 (PDB code 9B36)^27^, BPAM344-bound (without orthosteric ligands) inactive GluK2 (PDB code 8FWS)^32^, and glutamate-bound deep desensitized GluK2 (PDB code 9B38)^27^ structures. **(E, F)** Two designed disulfide crosslinks at the inter-protomer interface, shown in the homology model and the schematic figure.

**Figure S2.** Channel properties of the single cysteine mutant and protein preparation. **(A)** Representative electrophysiological traces from HEK293T cells expressing GluK2 WT, GluK2 K676C, and GluK2 N802C, activated by 10 mM glutamate. **(B)** Size exclusion chromatography profile of the GluK2 N802C/K676C mutant eluted from the Superose 6 column chromatography. The fractions used for the cryo-EM studies are indicated by the black dotted line. **(C)** SDS-PAGE analysis of the purified GluK2 N802C/K676C mutant, with the black line indicating the full-length GluK2 protein band.

**Figure S3.** Cryo-EM data processing workflow for structures of the GluK2 K676C/N802C mutant in the presence of glutamate and BPAM344 (a), and glutamate alone (b)

**Figure S4.** Cryo-EM analysis of GluK2 K676C/N802C. **(A, D)** Local resolution maps, **(B, E)** Fourier shell correlation (FSC) curves. The threshold was set at 0.143 for unmasked and masked curves, and a 0.5 threshold was used for the map-to-model curve. **(C, F)** Angular distribution of particles calculated using the 3D refinement reconstruction algorithm in cryoSPARC.

**Figure S5.** Comparison of ATD-LBD layers. **(A)** GluK2 structures, aligned parallel to the membrane. The distances between the center-of-mass (COM, black circles) of the ATD and LBD layers are indicated. **(B)** Superimposition of the ATD layer in the non-desensitized, intermediate, and desensitized GluK2 structures. The color codes match those in panel A. **(C)** ATD-LBD rotation. The LBD layers in the BPAM344- and glutamate-bound non-desensitized, and the BPAM344-, ConA-, and glutamate-bound open GluK2 conformations, are rotated counterclockwise when the ATD layers are superimposed.

**Figure S6.** Models and cryo-EM densities of the transmembrane domain. TM3 helices in non-desensitized, shallow-desensitized, intermediate, and deep-desensitized structures, with the protein structural model shown as ribbons and sticks. Cryo-EM densities are displayed in transparent gray.

## Notes

### Competing Interest Statement

The authors have declared no competing interest.

### Summary of Updates

Figure 5 revised; Section on discussion updated

