## Supplemental figures for "Structural Insights into Kainate Receptor Desensitization"

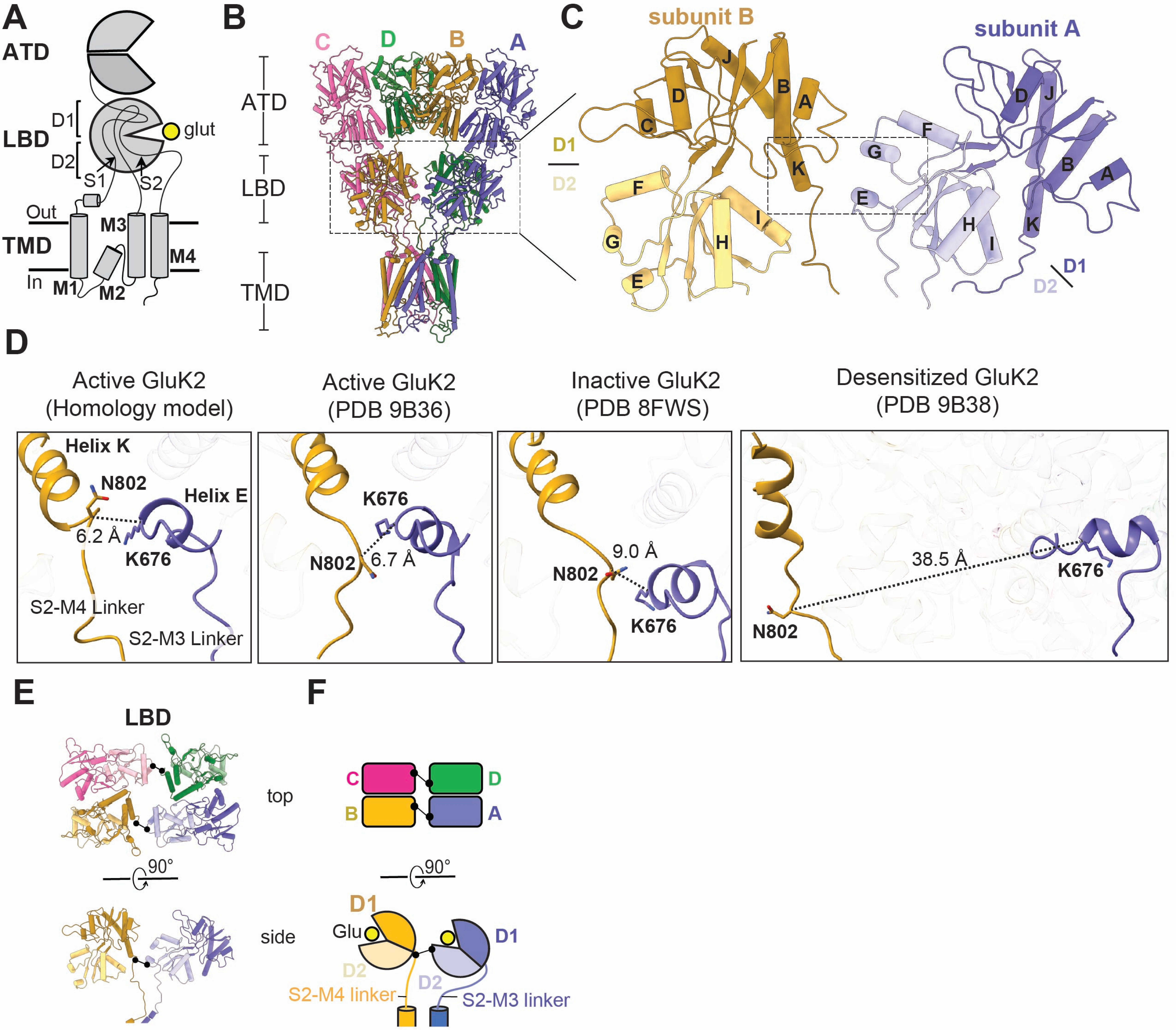

A
ATD
D1
LBD
glut
D2
S1
S2
Out
M3
TMD
M4
In
M1
M2
B
C D B A
ATD
LBD
TMD
C
subunit B
subunit A
D1
D2
D
Active GluK2
(Homology model)
Active GluK2
(PDB 9B36)
Inactive GluK2
(PDB 8FWS)
Desensitized GluK2
(PDB 9B38)
Helix K
N802
Helix E
6.2 Å
K676
S2-M4 Linker
S2-M3 Linker
6.7 Å
9.0 Å
38.5 Å
E
LBD
top
90°
side
F
C D
B A
D1
Glu
D2
S2-M4 linker
S2-M3 linker


### Figure S2

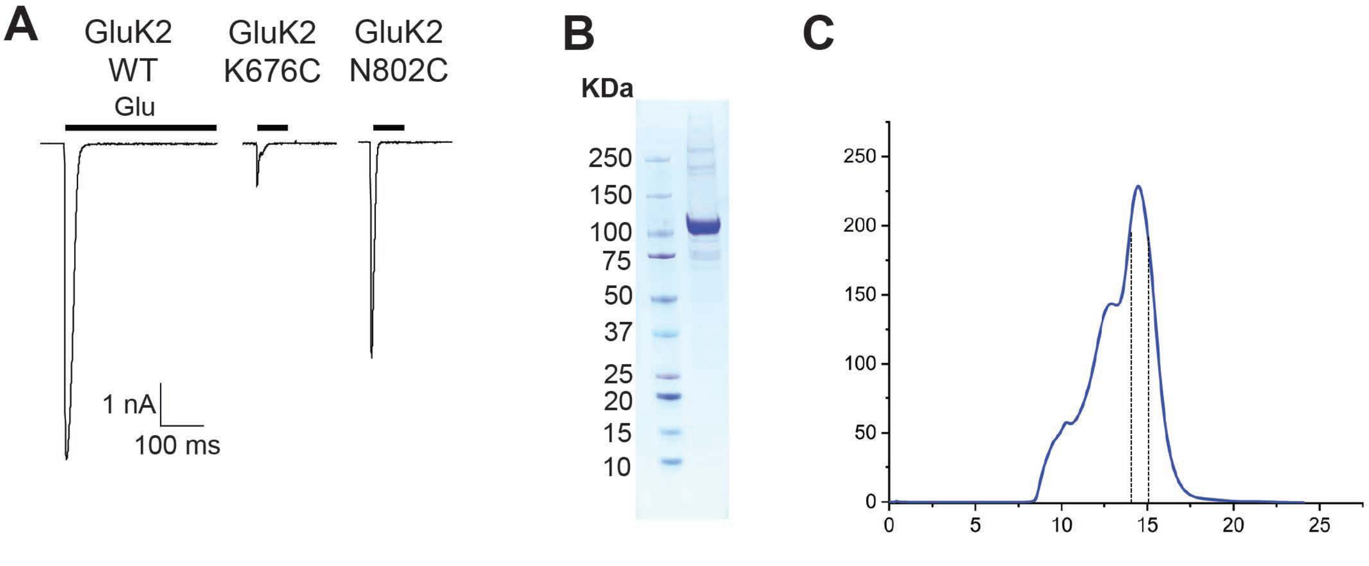
A
GluK2
WT
GluK2
K676C
GluK2
N802C
Glu
1 nA
100 ms
B
KDa
250
150
100
75
50
37
25
20
15
10
C
250
200
150
100
50
0
0
5
10
15
20
25

### Figure S2

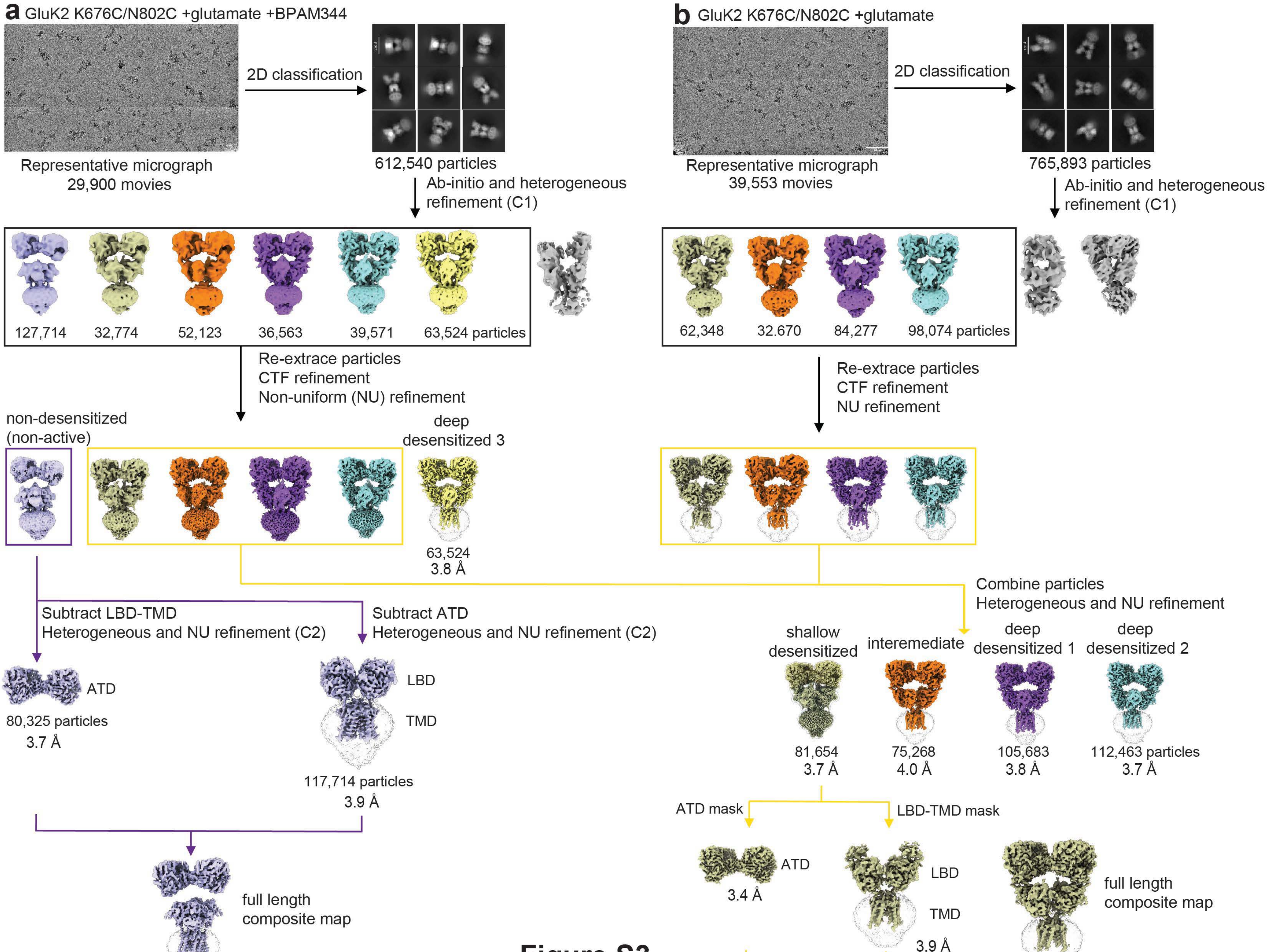
a GluK2 K676C/N802C +glutamate +BPAM344
2D classification
Representative micrograph
29,900 movies
612,540 particles
Ab-initio and heterogeneous refinement (C1)
127,714
32,774
52,123
36,563
39,571
63,524 particles
Re-extrace particles
CTF refinement
Non-uniform (NU) refinement
non-desensitized (non-active)
deep desensitized 3
63,524
3.8 Å
Subtract LBD-TMD
Heterogeneous and NU refinement (C2)
Subtract ATD
Heterogeneous and NU refinement (C2)
ATD
80,325 particles
3.7 Å
LBD
TMD
117,714 particles
3.9 Å
full length composite map
b GluK2 K676C/N802C +glutamate
2D classification
Representative micrograph
39,553 movies
765,893 particles
Ab-initio and heterogeneous refinement (C1)
62,348
32.670
84,277
98,074 particles
Re-extrace particles
CTF refinement
NU refinement
Combine particles
Heterogeneous and NU refinement
shallow desensitized
interemediate
deep desensitized 1
deep desensitized 2
81,654
3.7 Å
75,268
4.0 Å
105,683
3.8 Å
112,463 particles
3.7 Å
ATD mask
LBD-TMD mask
ATD
3.4 Å
LBD
TMD
3.9 Å
full length composite map

**Figure S3**

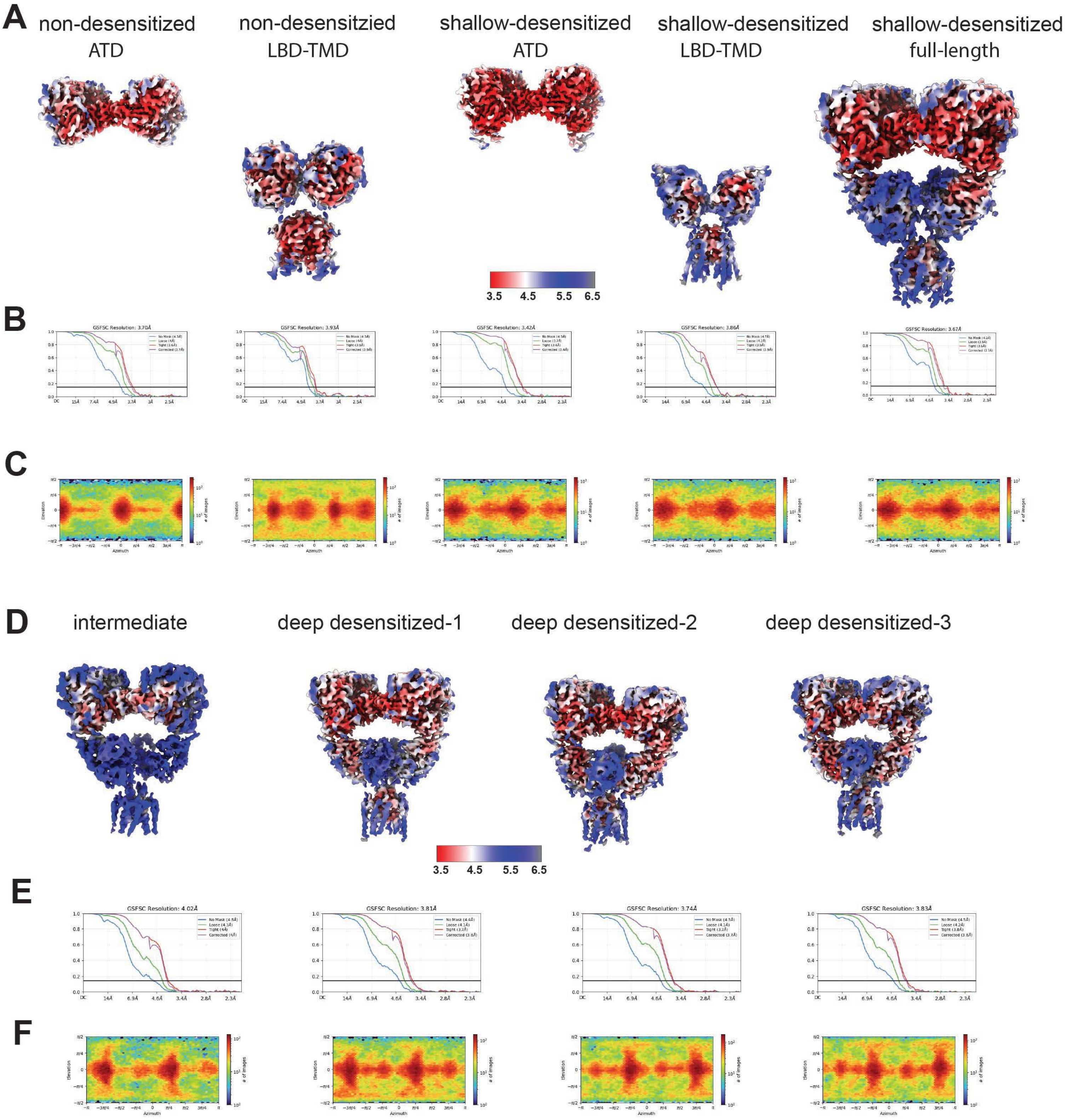
A
non-desensitized ATD
non-desensitzied LBD-TMD
shallow-desensitized ATD
shallow-desensitized LBD-TMD
shallow-desensitized full-length
3.5 4.5 5.5 6.5
B
GSFSC Resolution: 3.70Å
GSFSC Resolution: 3.93Å
GSFSC Resolution: 3.42Å
GSFSC Resolution: 3.86Å
GSFSC Resolution: 3.67Å
C
D
intermediate
deep desensitized-1
deep desensitized-2
deep desensitized-3
3.5 4.5 5.5 6.5
E
GSFSC Resolution: 4.02Å
GSFSC Resolution: 3.81Å
GSFSC Resolution: 3.74Å
GSFSC Resolution: 3.83Å
F

**Figure S4**

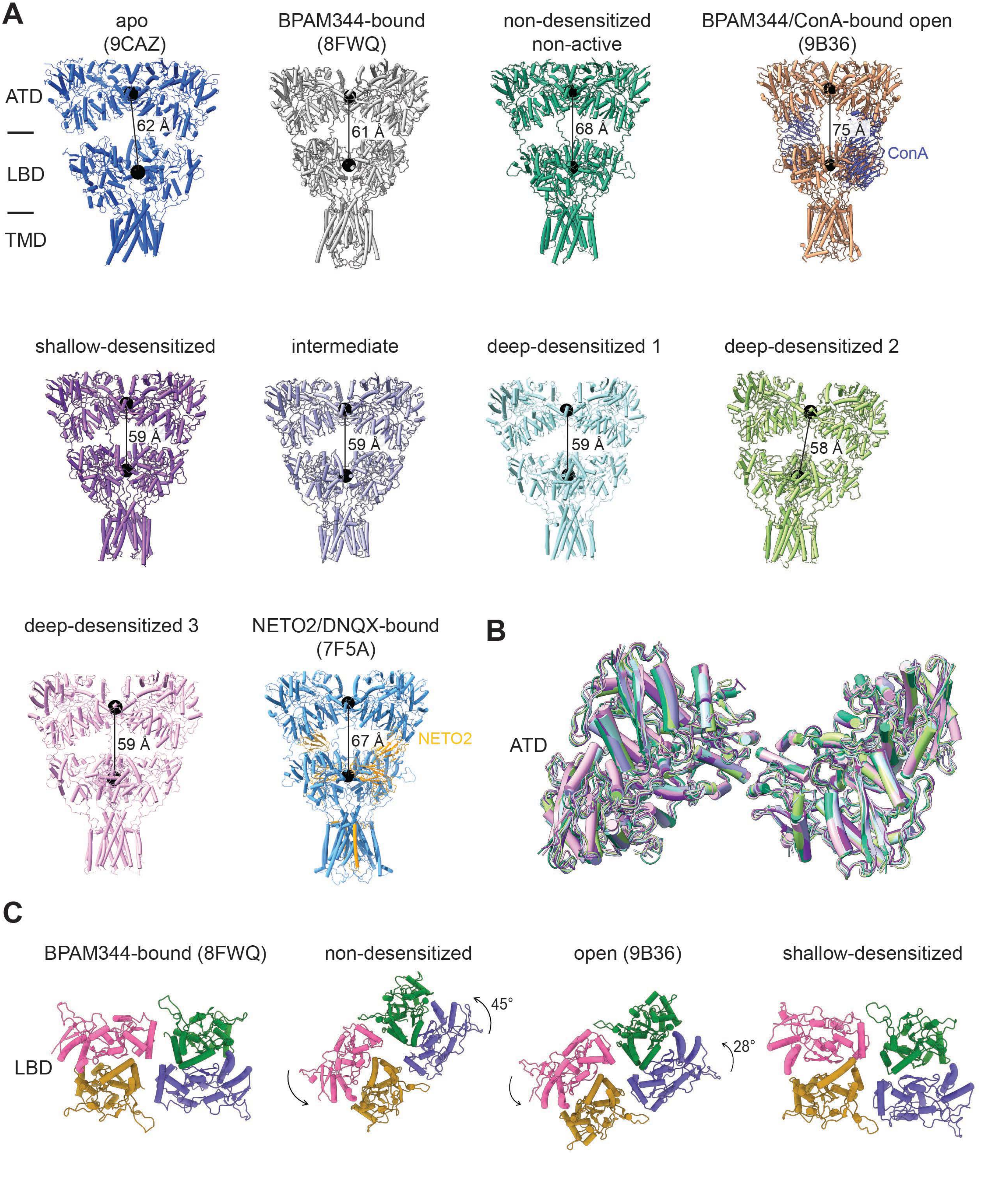
A
apo (9CAZ)
BPAM344-bound (8FWQ)
non-desensitized non-active
BPAM344/ConA-bound open (9B36)
ATD
LBD
TMD
62 Å
61 Å
68 Å
75 Å
ConA
shallow-desensitized
intermediate
deep-desensitized 1
deep-desensitized 2
59 Å
59 Å
59 Å
58 Å
deep-desensitized 3
NETO2/DNQX-bound (7F5A)
59 Å
67 Å
NETO2
B
ATD
C
BPAM344-bound (8FWQ)
non-desensitized
open (9B36)
shallow-desensitized
LBD
45°
28°

**Figure S5**

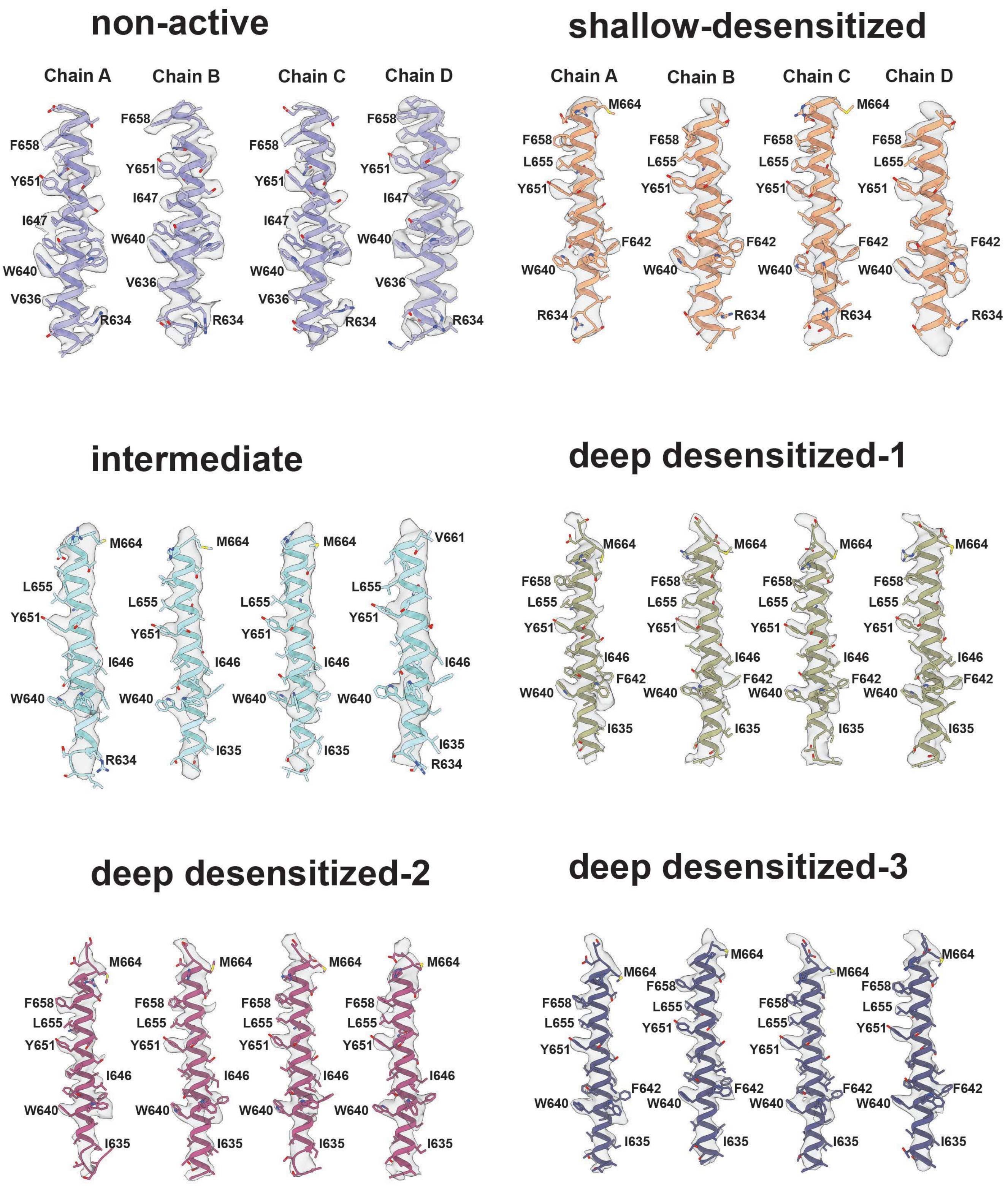
non-active
Chain A
Chain B
Chain C
Chain D
F658
Y651
I647
W640
V636
R634
shallow-desensitized
Chain A
Chain B
Chain C
Chain D
M664
F658
L655
Y651
F642
W640
R634
intermediate
M664
V661
L655
Y651
I646
W640
I635
R634
deep desensitized-1
M664
F658
L655
Y651
I646
F642
W640
I635
deep desensitized-2
M664
F658
L655
Y651
I646
W640
I635
deep desensitized-3
M664
F658
L655
Y651
F642
W640
I635

Figure S6

**Table S1. Cryo-EM data collection, refinement, and validation statistics.**

| | Non-desensitized ATD (EMD-48765) 9MZQ | Non-desensitized LBD-TMD (EMD-48766) 9MZR | Non-desensitized composite full-length (EMD-48767) 9MZS | Shallow-desensitized ATD (EMD-48762) 9MZN | Shallow-desensitized LBD-TMD (EMD-48763) 9MZO | Shallow-desensitized consensus full-length (EMD-48761) 9MZM | Shallow-desensitized composite full-length (EMD-48764) 9MZP |
|---|---|---|---|---|---|---|---|
| **Data collection and professing** | | | | | | | |
| Microscope | Titan Krios | Titan Krios | Titan Krios | Titan Krios | Titan Krios | Titan Krios | Titan Krios |
| Magnification (kx) | 81 | 81 | 81 | 81 | 81 | 81 | 81 |
| Voltage (kV) | 300 | 300 | 300 | 300 | 300 | 300 | 300 |
| Energy filter | Gatan | Gatan | Gatan | Gatan | Gatan | Gatan | Gatan |
| Energy filter slit width (eV) | 20 | 20 | 20 | 20 | 20 | 20 | 20 |
| Collection software | EPU | EPU | EPU | EPU | EPU | EPU | EPU |
| Electron exposure (e–/Å2) | 50 | 50 | 50 | 50 | 50 | 50 | 50 |
| Exposure rate (e–/Å2/frame) | 1 | 1 | 1 | 1 | 1 | 1 | 1 |
| Defocus range (μm) | -0.8 ~ -1.6 | -0.8 ~ -1.6 | -0.8 ~ -1.6 | -0.8 ~ -1.6 | -0.8 ~ -1.6 | -0.8 ~ -1.6 | -0.8 ~ -1.6 |
| Pixel size (Å) | 1.06 | 1.06 | 1.06 | 1.06 | 1.06 | 1.06 | 1.06 |
| Symmetry imposed | C2 | C2 | | C2 | C2 | C2 | |
| Initial particle images (no.) | 127,714 | 127,714 | 127,714 | 95,122 | 95,122 | 95.122 | 95,122 |
| Final particle images (no.) | 80,325 | 117,714 | 127,714 | 81.654 | 81,654 | 81.654 | 81.654 |
| 0.143 FSC half map masked (Å) | 3.78 | 3.96 | | 3.56 | 4.09 | 3.79 | |
| 0.143 FSC half map unmasked(Å) | 3.97 | 4.05 | | 3.71 | 4.19 | 3.96 | |
| **Refinement** | | | | | | | |
| Refinement package | Phenix | Phenix | Phenix | Phenix | Phenix | Phenix | Phenix |
| Initial model used (PDB code) | 9B36 | 9B36 | 9B36 | 9B36 | 9B36 | 9B36 | 9B36 |
| 0.5 FSC model resolution masked (Å) | 3.87 | 4.19 | 4.19 | 3.7 | 4.23 | 4.14 | 3.83 |
| 0.5 FSC model resolution unmasked (Å) | 4.13 | 4.28 | 4.29 | 3.84 | 4.32 | 4.22 | 4.03 |
| Map sharpening B factor ($Å^2$) | -143 | -163 | | -108 | -141 | -117.6 | |
| Model composition | | | | | | | |
| Non-hydrogen atoms | 12,582 | 11,927 | 25,004 | 12,784 | 11,857 | 24,825 | 24,825 |
| Protein residues | 1,534 | 1,501 | 3,093 | 1,565 | 1,489 | 3084 | 3,084 |
| Ligand | 23 | 4 | 29 | 24 | 9 | 30 | 30 |
| B-factors ($A^2$) | | | | | | | |
| Protein | 150.98 | 94.62 | 246.86 | 149.07 | 224.77 | 105.58 | 131.20 |
| Ligand | 173.91 | 35.82 | 240.10 | 144.34 | 188.59 | 103.13 | 135.65 |

| | | | | | | | |
|---|---|---|---|---|---|---|---|
| R.m.s. deviations | | | | | | | |
| Bond lengths (Å) | 0.004 | 0.003 | 0.002 | 0.003 | 0.004 | 0.003 | 0.002 |
| Bond angles (°) | 0.580 | 0.590 | 0.586 | 0.602 | 0.636 | 0.572 | 0.551 |
| Validation | | | | | | | |
| MolProbity score | 1.99 | 2.12 | 1.94 | 1.94 | 2.13 | 1.95 | 1.92 |
| Clashscore | 11.56 | 13.59 | 11.72 | 9.95 | 15.10 | 11.90 | 11.27 |
| Rotamer outliers (%) | 0 | 0.62 | 0.22 | 0.15 | 0.16 | 0.07 | 0 |
| Ramachandran plot | | | | | | | |
| Favored (%) | 93.84 | 92.30 | 94.83 | 93.64 | 93.08 | 94.85 | 94.88 |
| Allowed (%) | 6.16 | 7.70 | 5.17 | 6.36 | 6.92 | 5.15 | 5.12 |
| Disallowed (%) | 0 | 0 | 0 | 0 | 0 | 0 | 0 |
| CaBLAM outliers (%) | 3.49 | 3.07 | 2.52 | 2.65 | 3.50 | 3.18 | 3.41 |

| | intermediate (EMD-48760) 9MZL | deep desensitized 1 (EMD-48759) 9MZK | deep desensitized 2 (EMD-48758) 9MZJ | deep desensitized 3 (EMD-48757) 9MZI |
|---|---|---|---|---|
| **Data collection and professing** | | | | |
| Microscope | Titan Krios | Titan Krios | Titan Krios | Titan Krios |
| Magnification (kx) | 81000X | 81000X | 81000X | 81000X |
| Voltage (kV) | Gatan | Gatan | Gatan | Gatan |
| Energy filter | 20 | 20 | 20 | 20 |
| Energy filter slit width (eV) | EPU | EPU | EPU | EPU |
| Collection software | 300 | 300 | 300 | 300 |
| Electron exposure (e–/Å2) | 50 | 50 | 50 | 50 |
| Exposure rate (e–/Å2/frame) | 1 | 1 | 1 | 1 |
| Defocus range (μm) | -0.8 ~ -1.6 | -0.8 ~ -1.6 | -0.8 ~ -1.6 | -0.8 ~ -1.6 |
| Pixel size (Å) | 1.06 | 1.06 | 1.06 | 1.06 |
| Symmetry imposed | C1 | C1 | C1 | C1 |
| Initial particle images (no.) | 96,3254 | 129,840 | 107,645 | 70,524 |
| Final particle images (no.) | 84793 | 120,840 | 97,645 | 63,524 |
| 0.143 FSC half map masked (Å) | 4.16 | 3.94 | 3.89 | 4.26 |
| 0.143 FSC half map unmasked(Å) | 4.31 | 4.16 | 4.09 | 4.20 |
| **Refinement** | | | | |
| Refinement package | Phenix | Phenix | Phenix | Phenix |
| Initial model used (PDB code) | 9B39 | 9B39 | 9B39 | 9B39 |
| 0.5 FSC model resolution masked (Å) | 4.28 | 4.13 | 4.06 | 4.14 |
| 0.5 FSC model resolution unmasked (Å) | 4.40 | 4.28 | 4.24 | 4.29 |
| Map sharpening B factor ($Å^2$) | -102 | -104 | -105 | -107 |
| Model composition | | | | |
| Non-hydrogen atoms | 23,785 | 24,372 | 24568 | 24,582 |
| Protein residues | 2,989 | 3,008 | 3,036 | 3,037 |
| Ligand | 11 | 45 | 44 | 44 |
| B-factors ($A^2$) | | | | |
| Protein | 340.07 | 207.20 | 203.98 | 235.15 |
| Ligand | 239.79 | 199.40 | 233.78 | 228.36 |
| R.m.s. deviations | | | | |
| Bond lengths (Å) | 0.003 | 0.003 | 0.003 | 0.003 |
| Bond angles (°) | 0.638 | 0.620 | 0.598 | 0.621 |

| | | | | |
|---|---|---|---|---|
| Validation | | | | |
| MolProbity score | 2.15 | 2.06 | 2.05 | 2.09 |
| Clashscore | 16.55 | 13.56 | 13.31 | 15.75 |
| Rotamer outliers (%) | 0.15 | 0 | 0.15 | 0.04 |
| Ramachandran plot | | | | |
| Favored (%) | 93.43 | 93.71 | 93.61 | 94.24 |
| Allowed (%) | 6.54 | 6.29 | 6.36 | 5.76 |
| Disallowed (%) | 0.03 | 0 | 0.03 | 0 |
| CaBLAM outliers (%) | 4.07 | 3.44 | 3.26 | 3.13 |
